# Modulation of hepatic transcription factor EB activity during cold exposure uncovers direct regulation of bis(monoacylglycero)phosphate lipids by *Pla2g15*

**DOI:** 10.1101/2023.11.03.565498

**Authors:** Jessica W. Davidson, Raghav Jain, Thomas Kizzar, Gisela Geoghegan, Daniel J Nesbitt, Amy Cavanagh, Akira Abe, Kwamina Nyame, Andrea Hunger, Xiaojuan Chao, Isabella James, Helaina Von Bank, Dominique A. Baldwin, Gina Wade, Sylwia Michorowska, Rakesh Verma, Kathryn Scheuler, Vania Hinkovska-Galcheva, Evgenia Shishkova, Wen-Xing Ding, Joshua J Coon, James A. Shayman, Monther Abu-Remaileh, Judith A. Simcox

## Abstract

Cold exposure is a selective environmental stress that elicits a rapid metabolic shift to maintain energy homeostasis. In response to cold exposure, the liver rewires the metabolic state shifting from glucose to lipid catabolism. By probing the liver lipids in cold exposure, we observed that the lysosomal bis(monoacylglycero)phosphate (BMP) lipids were rapidly increased during cold exposure. BMP lipid changes occurred independently of lysosomal abundance but were dependent on the lysosomal transcriptional regulator transcription factor EB (TFEB). Knockdown of TFEB in hepatocytes decreased BMP lipid levels and led to cold intolerance in mice. We assessed TFEB binding sites of lysosomal genes and determined that the phospholipase Pla2g15 regulates BMP lipid catabolism. Knockdown of Pla2g15 in mice increased BMP lipid levels, ablated the cold-induced rise, and improved cold tolerance. Knockout of Pla2g15 in mice and hepatocytes led to increased BMP lipid levels, that were decreased with re-expression of Pla2g15. Mutation of the catalytic site of Pla2g15 ablated the BMP lipid breakdown. Together, our studies uncover TFEB regulation of BMP lipids through Pla2g15 catabolism.

## Introduction

The liver maintains energy homeostasis in mammals by acting as a metabolic transistor switching between providing glucose or lipids for peripheral tissue catabolism. In the fed state, the liver is a sink for excess glucose to maintain normoglycemia. When nutrients are scarce in fasting, the liver processes and packages lipids to fuel peripheral tissues and reserve glucose for the brain. The shift to systemic lipid utilization is a coordinated response between multiple organs that begins with white adipose lipolysis of triglyceride stores which releases free fatty acids into circulation, a portion of which are taken up by the liver and processed(Bornstein et al., 2023; Chitraju et al., 2020; Haemmerle et al., 2006; Schreiber et al., 2017). Understanding nutrient regulation in the liver during stress adaptation provides insight into how these processes are disrupted in the metabolic syndrome(Jurado-Fasoli et al., 2024).

One metabolic stress that causes rapid liver lipid remodeling is cold exposure (Simcox et al., 2017). Cold exposure is a selective pressure that increases white adipose tissue lipolysis, with excess free fatty acids being taken up into the liver for processing into complex lipids; acute cold exposure leads to temporary hepatic steatosis(Sostre-Colón et al., 2021; Zhang et al., 2024). Loss of this hepatic lipid processing in cold exposure leads to cold intolerance(Simcox et al., 2017). To identify other liver lipid metabolism pathways altered in cold exposure, we used untargeted liquid chromatography-mass spectrometry (LC/MS) based lipidomics. We identified over two hundred liver lipid species that were significantly altered with cold exposure(Jain et al., 2022; Simcox et al., 2017). These lipids represented a variety of chemically distinct lipid classes including acylcarnitines, triglycerides, ceramides, and phospholipids. We also noted 94 spectral features from our LC/MS analysis that were not identified but had reproducible, temperature-dependent differences in abundance. We wanted to determine the identity of these features and explore their function in mediating nutrient regulation in the liver.

Through assessing spectral feature fragmentation, we determined that several of these cold induced spectral features were bis(monoacylglycero)phosphate (BMP) lipids and validated these observations with targeted methods. BMP lipids are localized to the lysosome, where they mediate lipid and protein degradation(Kirkegaard et al., 2010; Locatelli-Hoops et al., 2006; Rodriguez-Navarro et al., 2012). Accumulation of BMP lipids is characteristic of a number of lysosomal disorders, however, their regulation and function remain poorly understood (Akgoc et al., 2015; Medoh and Abu-Remaileh, 2024; Showalter et al., 2020). To determine if increased BMP lipids correlated with altered lysosomal function, we profiled lysosomes using imaging and biochemical assays. We observed that lysosomes were localized to lipid droplets during cold exposure and that transcription factor EB (TFEB), a regulator of autophagy and lysosomal biogenesis, exhibited a cold-induced activity shift to regulation of lipid metabolism. Liver knockdown of TFEB resulted in decreased BMP lipids and cold intolerance in mice. We determined that TFEB regulated BMP lipid levels through the lysosomal phospholipase A2 gXV (Pla2g15) in hepatocytes. Our findings shed new light on the processes that govern liver adaptation to acute cold and on the regulation of BMP lipids via TFEB and Pla2g15. These findings hold the potential to modulate BMP lipids for the treatment of lysosomal storage disorders and metabolic disease.

## Results

### Lysosomal BMP lipids are increased in the liver during cold exposure in mice

To identify uncharacterized lipids that regulate liver fuel switching, we housed mice at room temperature (RT; 22-24°C) or cold (4°C) for 6h with food removed at the start of the experiment. Liver lipids were extracted by an organic solvent mixture and untargeted lipidomics by LC/MS was used to identify 712 lipids (**Figure 1A & S1A**). We observed that over 8000 additional MS/MS spectra collected in the experiment remained unannotated (unidentified ‘features’). To determine if some of these features represented identifiable compounds important for survival in cold, we filtered the features to remove background ions and retained features that: 1) had a retention time characteristic of lipids (1.0-15min); 2) had a mass within the expected lipid range (400-1500m/z); and 3) were present in at least 80% of samples. This reduced the dataset to 1233 unidentified features (**Figure 1B**). Next, we selected unidentified spectral features based on shared fragmentation patterns as well as the magnitude of fold change and significance between RT and cold using a false-discovery rate threshold of *q*<0.05 (**Figure 1C**). Using the LIPIDMAPS database (https://www.lipidmaps.org), we determined that two of the most significantly increased features were bis(monoacylglycero)phosphate (BMP) lipids, a class of lysosomal lipids (**Figure S1B**).

**Figure 1.**
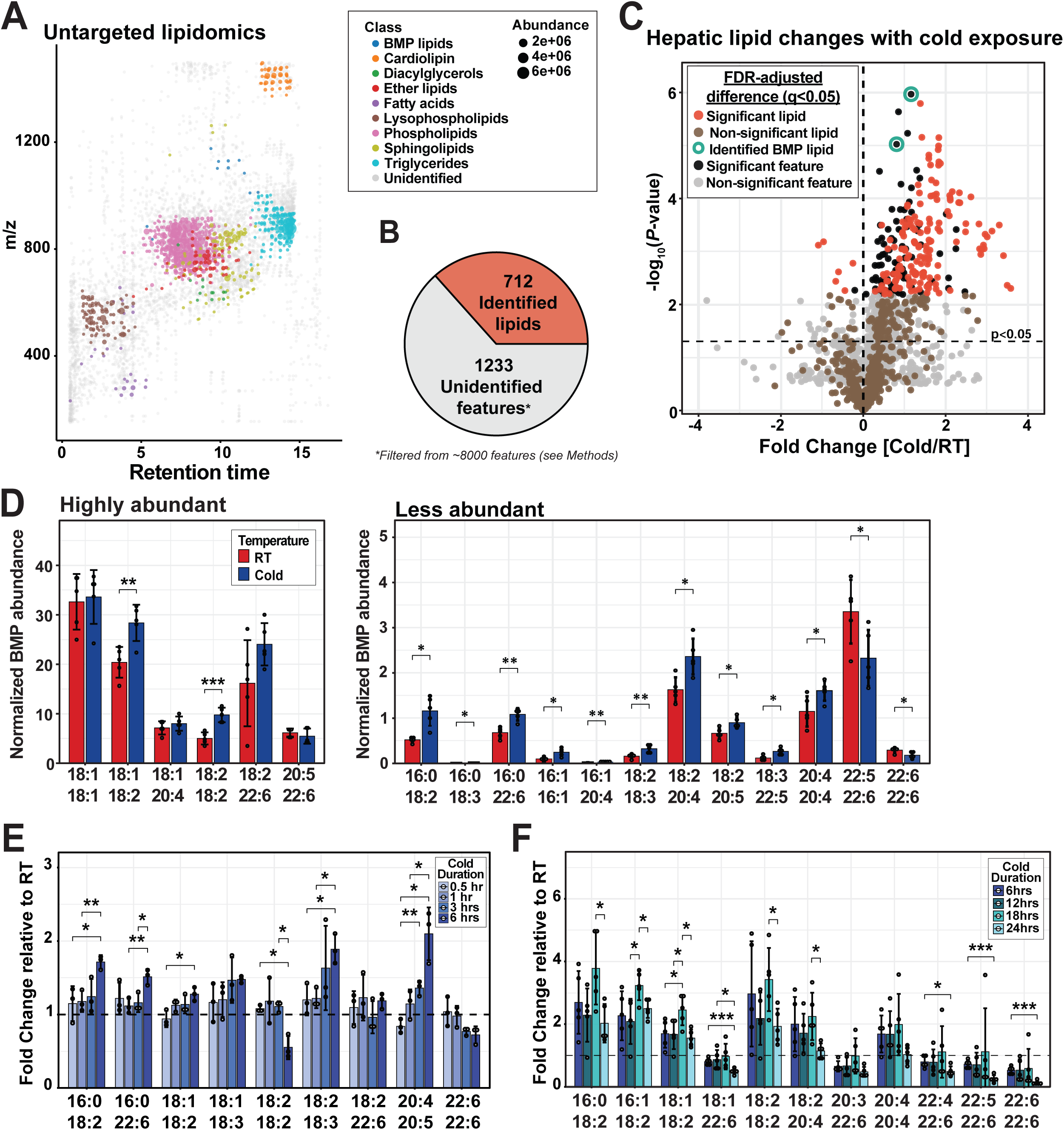
Lysosomal BMP lipids are increased in the liver during acute cold exposure in mice. A) Untargeted LC/MS analysis of mouse liver plotted to size (m/z) versus retention time with each dot representing a molecular feature. B) Pie chart representing the number of identified lipids and unidentified lipid-like features. Total unidentified features (∼8000) were filtered to remove background ions and contaminants and retain features present in at least 80% of samples. Remaining features were further filtered based on the retention time of lipid elution (0.5-15min) and mass-to-charge (m/z) range (200-1500m/z) of potential lipid species. C) Volcano plot of identified lipids and unidentified features (from Fig 1B) comparing RT (24°C) to cold (4°C) in the liver after 6h (n=6/group). Points in red or black represent significantly altered lipids or features, respectively, after false-discovery correction (*q*<0.05). Features circled in green were manually identified as lysosomal BMP lipid species based on the fragmentation pattern. D) Individual hepatic BMP lipid species were measured using a targeted LC/MS method in liver from RT and cold exposed mice (6 hour duration, n=5/group). The acyl chain composition of the BMP species are plotted on the x-axes. The most abundant BMP species are plotted on the left and significantly altered (*p*<0.05), less abundant BMP lipids are on the right. E) BMP lipid species changes in the liver during a short cold exposure time course of 0.5, 1, 3, and 6 hr, plotted as fold change relative to RT (n=3/group). F) BMP lipid species changes in the livers of mice cold exposed for up to 24 hours, reported as fold change relative to RT controls (n=5/group). Where relevant, mean±SD of experimental groups is plotted. BMP lipids abundances were normalized to internal standards and tissue weight. Student’s t-test was used for significance testing to compare lipid values. \**p*<0.05, \*\**p*<0.01, \*\*\**p*<0.001. Abbreviations: BMP, bis(monoacylglycero)phosphate; FDR, false discovery rate; LC/MS, liquid chromatography mass spectrometry; RT, room temperature.

To validate this finding, we developed a targeted LC/MS method to quantify 42 BMP lipid species and their precursor lipid phosphatidylglycerols (PGs)(Varadharajan et al., 2024). 14 individual BMP lipids were significantly increased in the liver during cold exposure with the majority containing essential fatty acids including 6 with an 18:2 acyl chain (**Figure 1D; Figure S1C**). The 18:2 fatty acid is a polyunsaturated fatty acid that is associated with increased membrane fluidity, potentially indicative of changes in BMP lipid function. We did not observe changes in liver PGs (**Figure S1D&E**). Interestingly, circulating BMP (*p*<0.05) and PG (*p*<0.01) lipids were significantly decreased during cold exposure (**Figure S1F**). Time course studies demonstrated that the rise in BMP lipids predominantly occurs in the liver at 6 hours (**Fig 1E**), but the elevation continues through a longer time course of 24 hours post cold exposure. (**Figure 1F & S1G**).

BMP lipids function as structural components of intralumenal vesicles within the endolysosomal system and are considered markers of lysosomal abundance. *In vitro* liposomal assays have demonstrated that polyunsaturation in BMP acyl chains facilitates BMP lipid interaction with lysosomal proteases and lipases, enhancing degradation of proteins and lipids in the lysosome lumen (Kirkegaard et al., 2010; Oninla et al., 2014).

The specific rise in polyunsaturated fatty acid containing BMP lipids could indicate a functional shift in lysosomes during cold exposure, or simply reflect a rise in lysosomal abundance.

### Lysosomes are localized to lipid droplets during cold exposure

To determine if the increase in BMP lipids with cold exposure was driven by increased lysosomal abundance, we assessed organelle abundance and localization in liver sections using electron microscopy (**Figure 2A**). There was no change in lysosomal abundance, however, lipid droplet abundance and lysosomal localization to lipid droplets was increased with cold exposure (**Figure 2B**). Other notable shifts in liver morphology included elongated mitochondria. We also quantified lysosomal (LAMP1, LAMP2, RAB5, and LIPA) and autophagy (LC3-I and LC3-II) protein levels which were unchanged, with the exception of LAMP1 which had a minor but significant increase with cold exposure (**Figure 2C**). Transcripts for lysosomal markers were largely unchanged between RT and cold except *Hexa* and *Atp6v1h* which were significantly increased with cold exposure, and *Lipa* and *Npc2* which were decreased with cold (**Figure 2D and S2A**).

**Figure 2.**
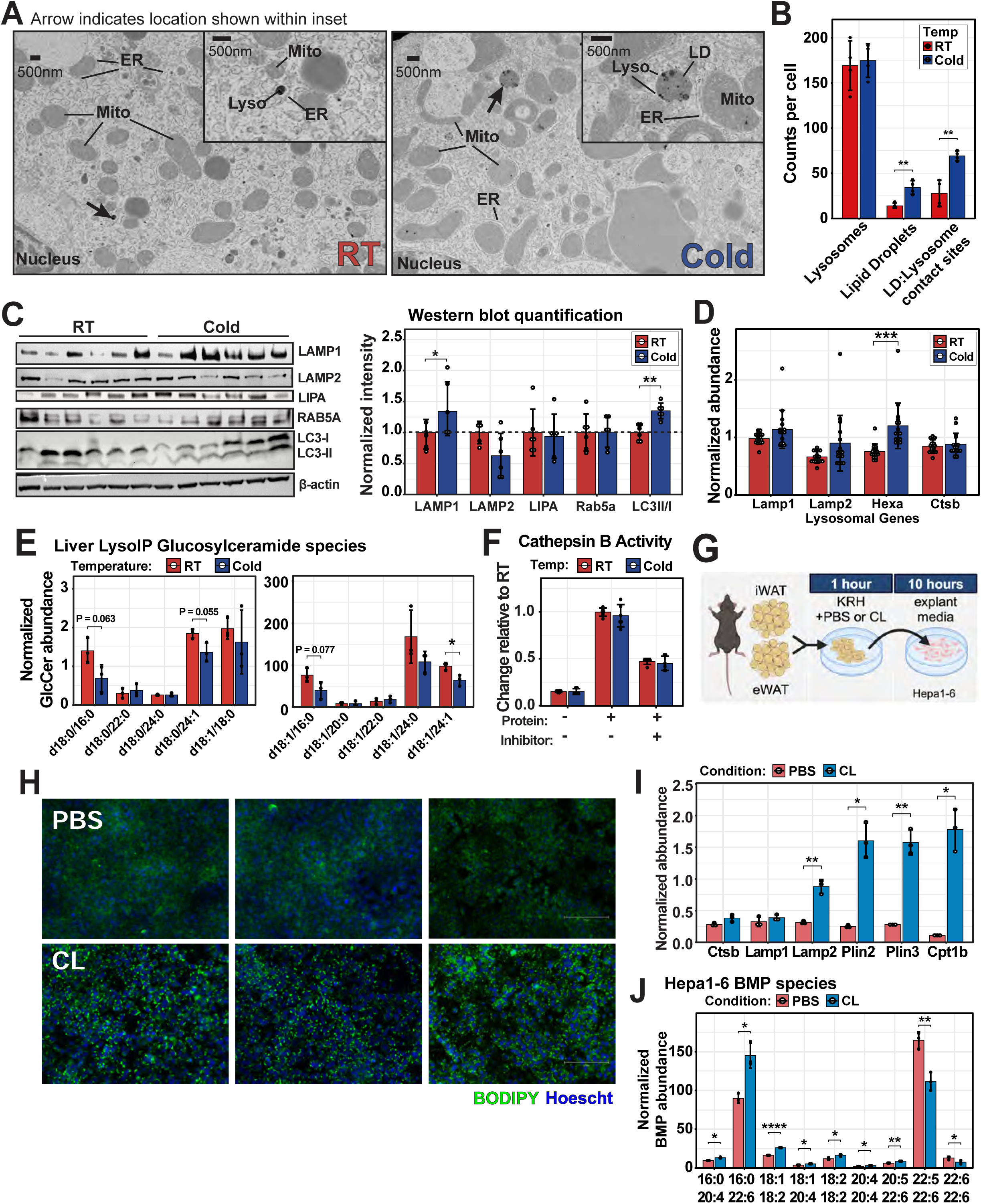
Lysosome localization is altered during cold exposure. A) Electron micrographs of liver sections from mice kept at either RT or cold exposure. Lysosomes are visible as dark, electron dense organelles. Arrows indicate zoomed in regions within insets. B) Quantification of lysosomes, lipid droplets (LD), and contact sites between lipid droplets and lysosomes. C) Western blot of lysosomal abundance and autophagy markers in liver from RT (24°C) and cold (4°C) exposed mice (n=6/group). Densitometry based quantification was normalized to average RT signal, and Wilcoxon non-parametric test used to calculate *P*-values. D) Lysosomal gene expression in liver from RT and cold exposed mice, assessed by qPCR (n=8/group; replicates shown). E) Glucosylceramide species in the lysosomes within the liver of RT and cold exposed LysoTag mice (n=3/group). F) Comparison of lysosomal cathepsin B protease activity in liver from mice at kept at RT or cold (n=6/group). G) Schematic of experimental design, mimicking cold exposure in Hepa 1-6 cells using adipose isolated from mice and treated with CL-316, 243. H) Bodipy and Hoescht staining of Hepa 1-6 cells incubated with explant media, in green and blue respectively. I) Gene expression of lysosomal and lipid droplet markers, as well as Cpt1b which is known to be induced by cold exposure in the liver (n=3/condition). J) Abundant BMP species that are significantly altered (*p*<0.05) in the CL condition are plotted (n=3/condition). All lipid measurements are normalized to internal standards. Where relevant, mean±SD of experimental groups is plotted. Student’s t-test was used for significance testing. \**p*<0.05, \*\**p*<0.01, \*\*\**p*<0.001, \*\*\*\**p*<0.0001. Abbreviations: CL, CL-316, 243.

To determine if the increased localization of lysosomes to lipid droplets coincided with a shift in function, we quantified lysosomal lipids in the liver using the LysoTag mouse (Laqtom et al., 2022) and cathepsin activity. Lysosomes are known to breakdown complex lipids including ceramides and glucosylceramides, with the activity of glucosylceramidases shifting in response to physiological stress such as fasting and high-fat diet(Longato et al., 2012; Xia et al., 2015). Aligned with these observations, there was a significant decrease in glucosylceramide d18:1/24:1 with cold exposure, and a trend in decreased d18:0/16:0, d18:0/24:1, and d18:1/16:0 glucosylceramides which would indicate increased glucosylceramidase activity (**Figure 2E**). The isolated lysosomes also had the expected cold-induced increase in BMP lipids (**Figure S2G**). There were no changes in Cathepsin B activity (**Figure 2E**). These results suggest a targeted shift in lysosomal activity, specifically towards lysosomal lipid processing.

One challenge in determining the mechanism of liver BMP lipid regulation in cold exposure is understanding the cells that contribute to the lipid phenotype. Recent work has shown that hepatocytes are the primary contributor to BMP lipids in the liver and to confirm this we used a culture model of hepatocytes (Hepa 1-6 cells) (Grabner et al., 2020). The hepatocyte response to cold exposure is mediated by free fatty acid lipolysis in white adipose tissue(Simcox et al., 2017). To recapitulate this lipolysis, we treated white adipose tissue (WAT) explants with β3-adrenergic receptor agonist CL-316,243 (CL) to induce lipolysis or vehicle control (PBS), and then transferred the media to the Hepa 1-6 cells (**Figure 2G**). We observed that Hepa 1-6 cells with CL explant treatment have increased lipid droplet abundance, similar to hepatocytes in the liver (**Figure 2B & 2H**). We observed increases in transcripts for lysosomal markers (*Lamp2*), lipid droplet markers (*Plin2, Plin3*), and established mitochondrial markers responsive to cold exposure (*Cpt1b*) (**Figure 2I**)(Simcox et al., 2017). We also observed a significant increase in specific BMP lipid species that corresponds to the changes we observed in the liver (**Figure 2J**). These results suggest that the major changes observed in the liver, are driven by hepatocyte specific differences in response to adipocyte lipolysis.

In total, biochemical analysis show increased lysosomal localization to lipid droplets with cold exposure and increased lysosomal lipid processing. These results also suggest that the differences in BMP lipids are driven by changes in hepatocytes, and that these changes are in response to altered white adipose tissue lipolysis. We next sought to determine the regulation of these changes in lysosomal function and localization.

### The master lysosomal regulator transcription factor EB is activated during cold exposure

We evaluated liver transcript levels for genes involved in lysosome biogenesis and function as well as related processes including autophagy (*Gabarapl1,Sqstm1*), mitochondrial oxidation (*Ppargc1a, Cpt1b*) (Jain et al., 2022; Simcox et al., 2017), endosome (*Ulk1, Stx17, Wdr45*), and lipid droplet formation (*Plin2, Dgat1, Pdzd8, Plin3*) (**Figure 3A & S2A-F**). There were coordinated changes including increases in autophagy makers *Gabarapl1* and *Sqstm1(Sardiello et al., 2009)*, and increases in lysosomal markers *Atp6v1h* and *Hexa*. Many of these transcripts contain a ‘CLEAR’ (<u>c</u>oordinated lysosome <u>e</u>xpression <u>a</u>nd regulation) sites upstream and downstream of their promoters (Palmieri et al., 2011; Sardiello et al., 2009). These sites are bound by transcription factors that are members of the MiTF (microphthalmia-associated transcription factor) family. The most abundant MiTF member in hepatocytes is transcription factor EB (TFEB)(Azimifar et al., 2014; Niu et al., 2022). There were also significant expression changes in *Tfeb* and *Tfe3* (**Figure S2B**). To determine if there was altered TFEB activity, we assessed liver protein levels of phosphorylated and unphosphorylated TFEB observing a decrease in phosphorylation (**Figure 3B**) coinciding with increased nuclear (active) TFEB with cold exposure compared to cytosolic (inactive) TFEB (**Figure 3C**). The nuclear-to-cytosolic ratio of TFEB was 3-fold higher in cold than RT (*p*<0.01; **Figure 3C**). Though most studies of TFEB activity characterization have focused on lysosomal activation, previous work has implicated TFEB in the control of hepatic lipid metabolism, including through activation of *Ppargc1a* **(Figure S2B)***(Gosis et al., 2022; Settembre et al., 2013)*.

**Figure 3.**
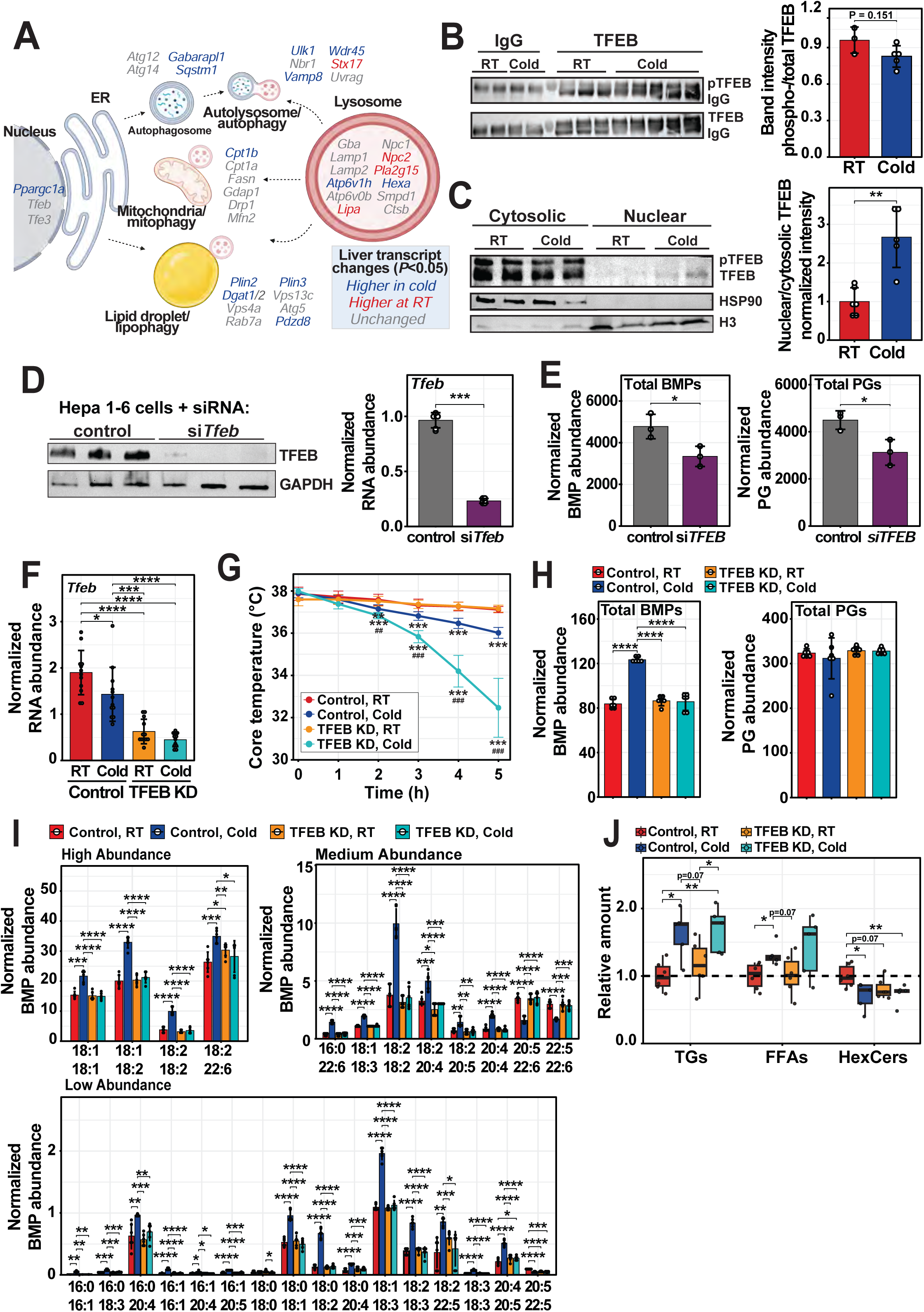
Hepatic knockdown of TFEB causes cold intolerance in mice. A) Schematic of potential lysosomal activities and associated genes for each process/organelle function. Genes are colored based on qPCR transcriptional changes (*p*<0.05) between RT and cold exposed murine livers (n=6/group). B) TFEB immunoprecipitation and western blot of TFEB phosphorylation in the liver of RT (n=3) and 6h cold exposed mice (n=5). IgG was used as a control for the immunoprecipitation. The upper bands, representing phospho-(top blot) and total (bottom blot), were quantified based on densitometry, and the ratio reported. C) Western blot of TFEB from liver of mice kept at RT or cold (n=6/group). Liver lysate was fractionated into nuclear and cytosolic fractions prior to western blot. Bands were quantified using densitometry and the ratio of nuclear (active) to cytosolic (inactive) TFEB signal was plotted. D) Western blot of TFEB protein in Hepa1-6 cells treated with siRNA targeting *Tfeb* or a scramble sequence as control is shown (n=3/condition). Gene expression of *Tfeb*, measured by qPCR, is to the right (n=3/ condition, replicates shown). E) Total BMP and PG lipids of Hepa1-6 murine hepatocytes treated with siRNA against *Tfeb* or a scramble sequence control is plotted (n=3/group). Lipids are normalized to internal standards and protein content. F) Hepatic *Tfeb* expression in mice treated with AAV8-GFP-U6-scrmb-shRNA (control; n=12) or AAV8-GFP-U6-TFEB-shRNA (TFEB knockdown (KD); n=12), and exposed to 5 hours of RT or cold three weeks after AAV injection. G) Core body temperature was measured hourly via rectal probe in control or TFEB KD mice kept at RT or cold for 5h (n=6/group). A two-way ANOVA with interaction was used to assess the effect of KD and environmental temperature on core body temperature. Asterisks (*) denote significant (*p*<0.05) difference between core temperature due to temperature of mice with the same KD status. Pound sign (#) indicates significant difference in core temperature between control and KD mice kept at 4°C. H) Total liver BMP lipids and PGs for each group from the cold tolerance test (n=6/group), assessed by targeted lipidomics. I) BMP lipid species significantly altered during cold exposure are plotted (n=6/group). Lipid abundances normalized to internal standards and tissue weight. J) Lipid abundance of various lipid classes in in the liver are measured by untargeted LC/MS lipidomics are plotted. Values are relative to RT controls for each lipid class. Where relevant, mean±SD of experimental groups is plotted. \**p*<0.05, \*\**p*<0.01, \*\*\**p*<0.001, \*\*\*\**p*<0.0001. Abbreviations: HexCer, hexosylceramides; BMP, bis(monoacylglycero)phosphate; Con, control; FFA, free fatty acids; KD, knockdown; LC/MS, liquid chromatography mass spectrometry; RT, room temperature; TFEB, transcription factor EB; TG, triglycerides.

To determine if *Tfeb* modulated BMP lipids in hepatocytes, we used siRNA to knock down (KD) *Tfeb* in Hepa1-6 cells (**Figure 3D**). Controls were treated with siRNA containing a scramble sequence. After 48h, cells were harvested for BMP analysis, total BMP lipids were significantly decreased in *Tfeb* KD cells as were PG levels (*p*<0.05; **Figure 3E-F**). The knockdown of TFEB in fed mice led to decreased BMP lipid levels (**Figure S3A**). These results identified TFEB as a novel genomic regulator of BMP lipids in hepatocytes. Given the increase in nuclear-localized TFEB and regulation of BMP lipids by TFEB in hepatocytes, we aimed to determine if TFEB was functionally important in cold tolerance and the hepatic lipid response.

### Hepatic TFEB is required for cold tolerance in mice

To assess the requirement of hepatic TFEB in the cold response, we knocked down *Tfeb* using shRNA encoded in an AAV8 vector which targets hepatocytes. After three weeks, control and KD mice were subjected to 5h at RT (23°C; n=5-6 and 6, respectively), or 5h in cold (4°C; n=5-6 and 6, respectively) with food removed at the start of the experiment. In the liver, KD mice had ∼70% lower *Tfeb* gene expression than controls (**Figure 3F)**. Mice kept at 4°C had significantly lower core body temperature than those at RT after 2h. Decreased body temperature in the cold was exacerbated with loss of TFEB (*p*<0.01 at all timepoints; **Figure 3G**). Loss of TFEB also ablated cold induced differences in BMP lipids (**Figure 3H&I**), but changes in other lipids persisted with loss of TFEB (**Figure 3J, S3C&G**). Despite the major shift in body temperature with loss of TFEB, there were no changes between control and KD in body weight loss or transcripts associated with lipid droplets and mitochondria (**Figure S3B-E**). KD of TFEB did lead to increased liver weight and shifts in lysosomal transcripts (**Figure S3 B,D, &F).** These results show that hepatic TFEB is required for acute cold adaptation in mice, and that this regulation leads to altered lysosomal transcripts and BMP lipids, without major changes in other hepatic or circulating lipids.

### Cold exposure rewires TFEB to coordinate lipid metabolic processes

Given the impact of liver TFEB loss on cold tolerance and BMP lipids, we sought to understand how TFEB regulates BMP lipids. Because TFEB is a transcriptional regulator, we performed ChIP-seq for TFEB in liver of mice housed at RT or fasted in cold for 6h (**Figure 4A&S4A**). After filtering reads using a stringent FDR, we observed that there was a drastic shift in the type of binding sites between conditions. The percentage of intronic binding sites (of total sites) decreased from 61.1% in RT to 48.4% in cold (*q*<0.05; **Figure 4B**). The majority of binding sites were for protein coding genes, and the greatest increase was for intergenic regions (RT 24.1% to cold 36.2%) while the percentage of promoter binding sites were similar in both conditions (RT 7.8 and cold 6.0%) (**Figure S4B)**. We validated these results observing TFEB binding to established targets such as *Ctsa*, *Ctsb*, and *Npc1* (**Figure 4C&S4C**)(Sardiello et al., 2009). This was confirmed using targeted ChIP-qPCR based on binding sites annotated to TFEB from sequencing data (**Figure 4C**). These differences in TFEB activity are potentially driven by a shift in TFEB binding partners as observed by LC/MS of TFEB co-immunoprecipitation (**Figure S4D**). The regulation by TFEB was confirmed by ablation of the cold-induced decrease in *Ctsa* and *Ctsb* expression in TFEB KD mouse livers (**Figure 4D**).

**Figure 4.**
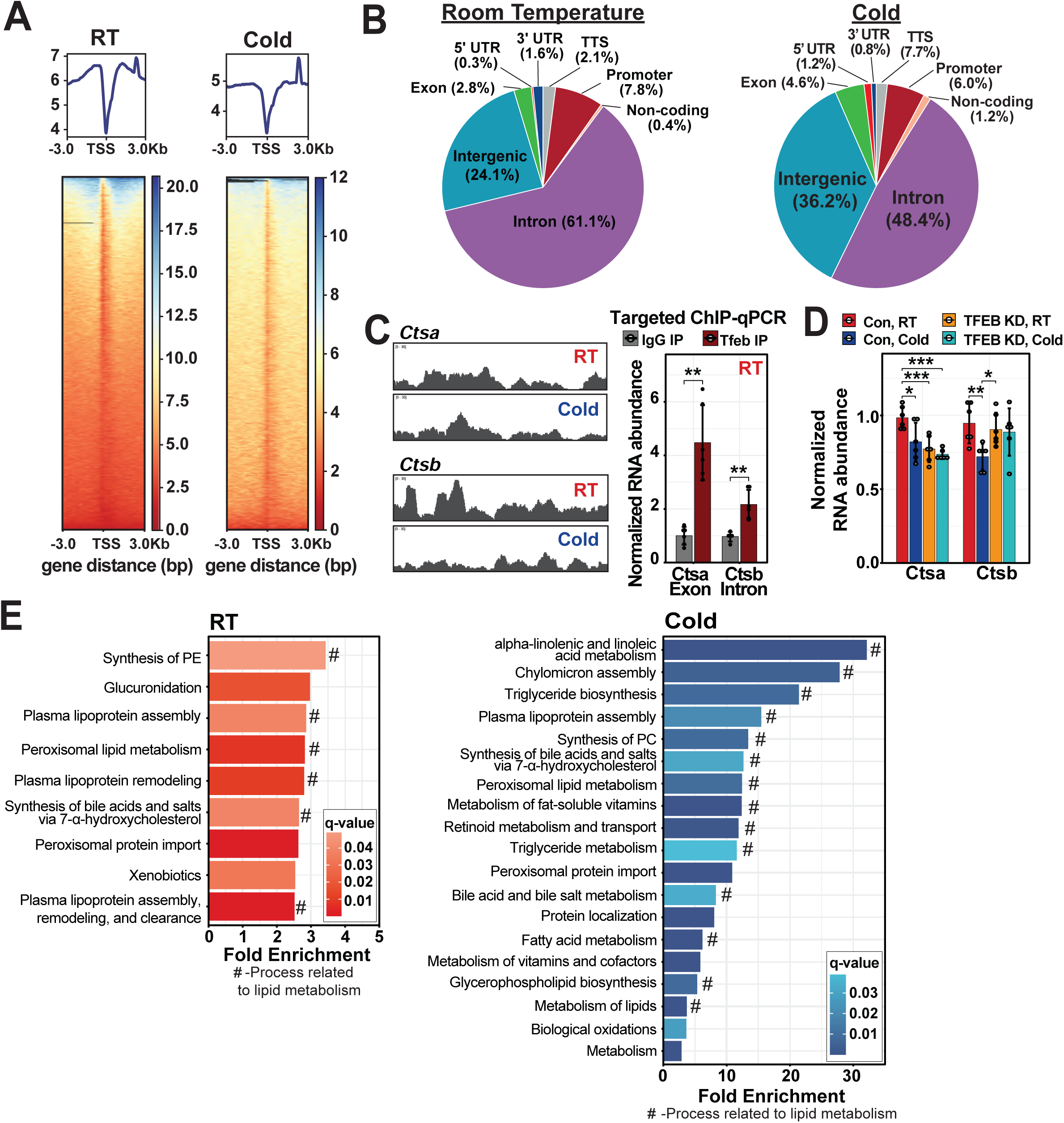
ChIP-seq of hepatic TFEB indicates a reprogramming towards lipid metabolism during cold exposure. A) Transcription start site-centered heat maps comparing hepatic TFEB binding between RT and cold based on ChIP-seq data. B) Pie chart describing the distribution of TFEB genomic binding sites during RT or cold. C) ChIP tracks of previously described genes known to be bound by TFEB are shown^13^. Targeted ChIP-qPCR was performed to validate binding sites at regions marked by red pound signs on the ChIP tracks. D) Gene expression of Ctsa and Ctsb in the liver of control and TFEB knockdown mice kept at RT or exposed to cold for 6 hours (n=6/group). E) Reactome of genes annotated to TFEB binding sites at RT or with 6 hours of cold exposure. Of processes represented in the gene sets, those related to lipid metabolism are annotated with a ‘#’. All ChIP analyses were performed after false-discovery correction with a *q*<0.05 cutoff. Where relevant, mean±SD of experimental groups is plotted. \**p*<0.05, \*\**p*<0.01, \*\*\**p*<0.001. Abbreviations: ChIP-seq, chromatin immunoprecipitation sequencing; RT, room temperature

To identify the processes coordinated by TFEB in cold, we performed ‘Reactome analysis’ using genes annotated to TFEB in Cold as input and compared them to RT. We observed that 75% of all reported Reactome processes related to lipid metabolism (**Figure 4E**). Many of the most significantly (*q*<0.05) enriched processes were 10-30 fold above expected representation in the gene set. With our findings from TFEB KD mice, these data indicated that TFEB is responsible for rewiring liver lipid metabolism that occurs in cold exposure.

### TFEB regulates BMP lipids via *Pla2g15* in hepatocytes *in vitro*

Due to the altered BMP lipid levels in TFEB KD hepatocytes, we postulated that TFEB regulates enzymes directly involved in the metabolism of BMP lipids in hepatocytes. To identify potential enzymatic BMP lipid regulators in the lysosome, we overlaid all unique genes identified in our TFEB ChIP-seq analysis with proteins localized to the lysosome(Abu-Remaileh et al., 2017; Eapen et al., 2021; Yu et al., 2024), and genes with expression significantly altered in cold exposure compared to RT after 6h (*q*<0.05; **Figure 5A**). We filtered the resulting list of 24 genes to 4 candidates that are predicted to be regulators of lipid metabolism (**Figure 5A**). Of these genes, only one had altered expression in cold and in TFEB knockdown cells, phospholipase A2 group XV (*Pla2g15*) (**Figure 5B &C**). Pla2g15 expression and protein levels were decreased with cold exposure (**Figure 5D**). There was a sharp peak upstream of the promoter of *Pla2g15* in both TFEB and RNA pol ChIP-seq – indicative of active transcription. Targeted ChIP confirmed TFEB binding to the Pla2g15 site (**Figure 5E**). There was also a CLEAR site present within the coding sequence 1543 base pairs downstream of the transcription start site (sequence: GTCACGTGTG)(Palmieri et al., 2011). Luciferase assays of the TFEB binding sites demonstrated TFEB occupancy of the Pla2g15 promoter, that was increased in TFEB overexpression (**Figure 5F & S5A**).

**Figure 5.**
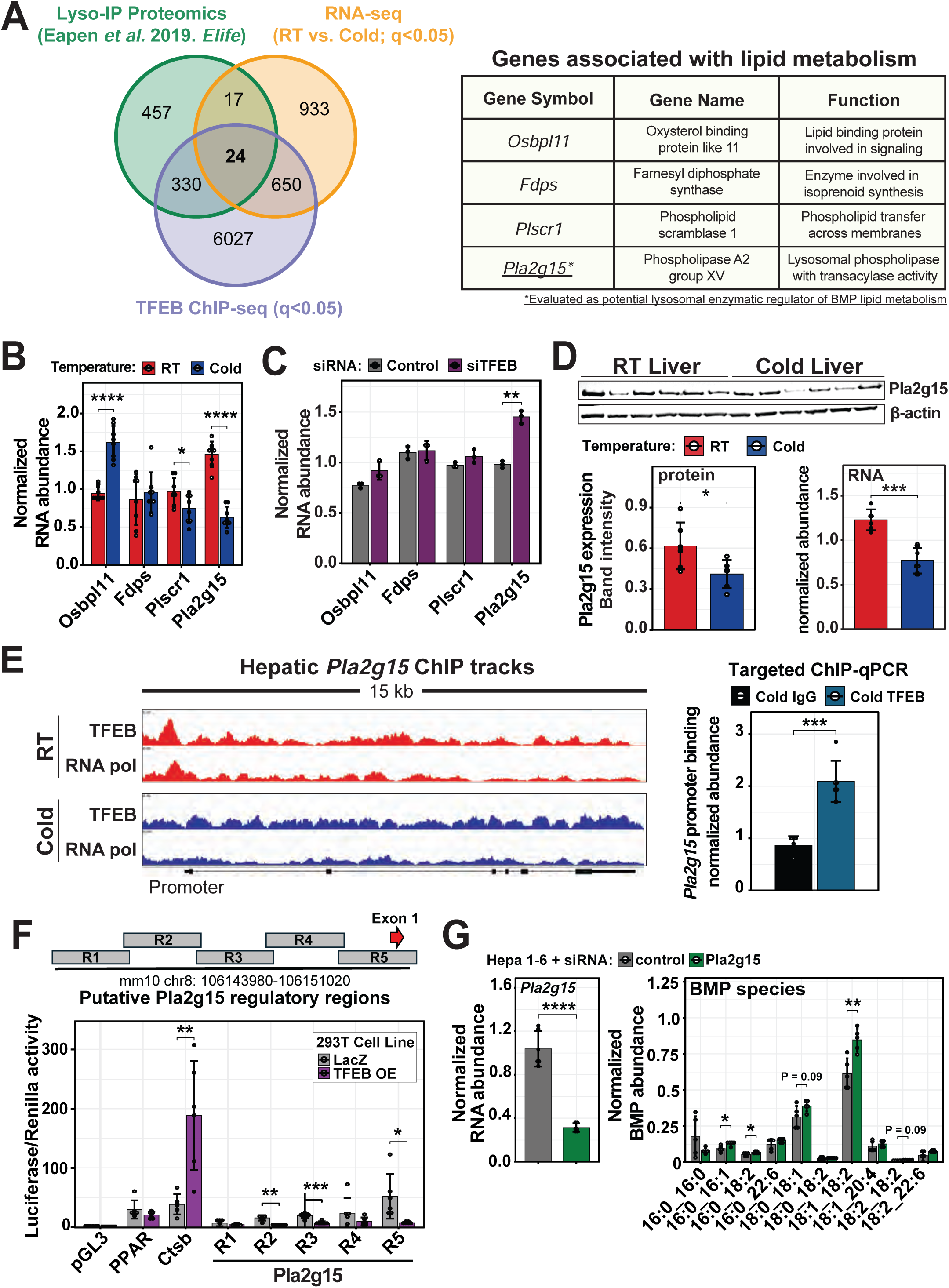
TFEB regulates BMP lipids in hepatocytes via Pla2g15 *in vitro*. A) Overlay of identified lysosomal proteins from Eapen *et al*. (2021)^26^ with all unique genes from TFEB ChIP-seq (*q*<0.05) and all genes with expression significantly altered between RT and cold after 6h (*q*<0.05). Of the 24 genes in the center overlap, four were annotated with lipid functions and are shown in the table to the right. B) Expression of the candidate genes in liver from mice kept at RT or cold exposed for 6 hours is shown (n=8/group). C) Candidate gene expression in Hepa1-6 cells treated with siRNA against TFEB or a scrambled control (n=3/group). D) Western blot of Pla2g15 is shown for RT and cold exposed mice (n=6/group). Band intensity, normalized to the loading control b-actin, is plotted. Gene expression, measured by qPCR, is also shown for *Pla2g15* in RT and cold exposed livers (n=6/group). E) ChIP tracks of TFEB and RNA polymerase on the genomic region encoding *Pla2g15* in RT and cold livers. To the right, targeted ChIP-qPCR is shown for the binding regions of TFEB to the promoter of *Pla2g15* in the cold. IgG was used as a control. F) Schematic of Pla2g15 regulatory regions expressed upstream of luciferase in LacZ or TFEB overexpressing 293T cells, in the luciferase binding assay below. Luciferase activity was normalized to Renilla as a control (n=6/condition, representative result from 3 independent experiments). G) Gene expression of *Pla2g15* in Hepa1-6 cells treated with siRNA targeting Tfeb or scramble sequence as control (n=3/group, 2 replicates per sample). Select BMP lipid species were measured from Hepa1-6 cells following treatment with siRNA against *Pla2g15* or a scramble sequence control (n=5/group). Lipids normalized to internal standards and protein. Mean±SD of experimental groups is plotted. \**p*<0.05, \*\**p*<0.01, \*\*\**p*<0.001, \*\*\*\**p*<0.0001. Abbreviations: BMP, bis(monoacylglycero)phosphate; TFEB, transcription factor EB.

To determine if Pla2g15 regulates BMP lipid levels, we performed a knockdown in Hepa 1-6 cells using siRNA (**Fig5G**). Knockdown of Pla2g15 led to an increase in BMP lipids including BMP 18:1_18:2, one of the primary BMP lipids increased *in vivo* with cold exposure. We also took an orthogonal approach and performed a KD of mannose-6-phosphate receptor (*M6pr*), which is required to traffic the Pla2g15 protein to lysosomes(Abe et al., 2008), and observed that total BMP lipids increased 2-fold with *M6pr* KD (*p*<0.05; **Figure S5B**).

Together, these data suggest that TFEB is a genomic regulator of *Pla2g15*, and that Pla2g15 may enzymatically regulate of BMP lipids. In this model, TFEB binding to the *Pla2g15* promoter is inhibitory and Pla2g15 degrades BMP lipids into lysophosphatidylglycerol and fatty acids (**Figure 6 A&B**). During cold exposure, increased TFEB binding of the *Pla2g15* promoter would decrease Pla2g15 expression, leading to the observed increase in BMP lipid levels.

**Figure 6:**
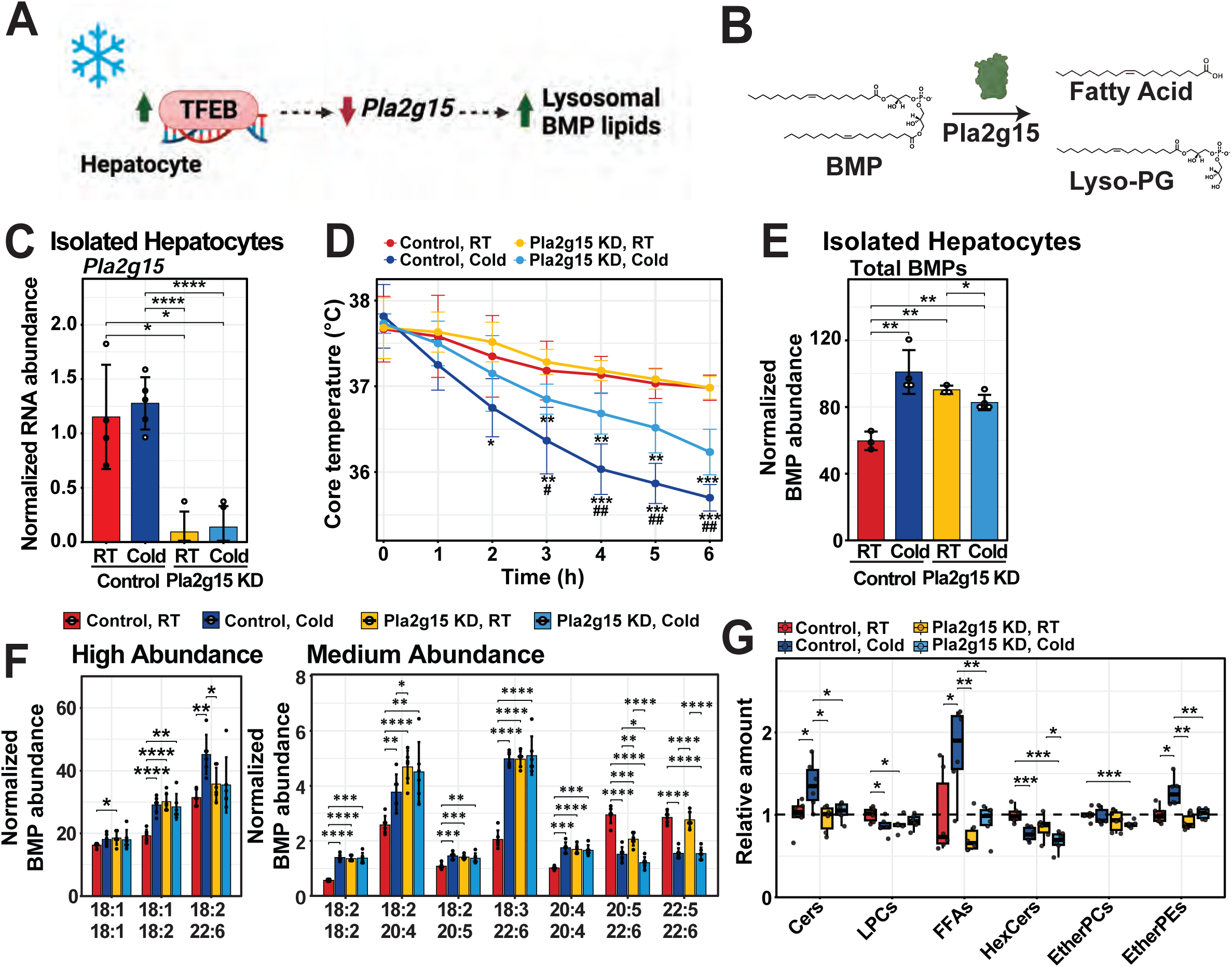
Pla2g15 is responsible for cold induced changes in BMP lipids. A) Model summarizing the regulation of BMP lipids by TFEB and *Pla2g15*. B) Proposed mechanism of Pla2g15 degradation of BMP lipids. C) *Pla2g15* gene expression in hepatocytes isolated from mice three weeks post treatment with AAV8-GFP-U6-scrmb-shRNA (control; n=9) or AAV8-GFP-U6-mPla2g15-shRNA (Pla2g15 knockdown (KD); n=9) and immediately following 6h at either RT (n=4/group) or cold (n=5/group). D) Results of a cold tolerance test in control and Pla2g15 KD mice kept at RT or in the cold for 6h, where core body temperature was recorded hourly (n=6/group). A two-way ANOVA with interaction was used to assess the effect of Pla2g15 KD and environmental temperature on core body temperature. Significant differences (p<0.05) in core temperature due to environmental temperature of mice within the same group (Pla2g15 KD or controls) are indicated by asterisks (*). Significant differences in core temperature between the control and KD mice in the cold (4°C) are indicated by a Pound sign (#). E) Total BMP lipids in hepatocytes isolated from control mice kept at RT or cold (n=3 and n=4, respectively) and Pla2g15 KD mice kept at RT and cold for 6hrs (n=3 and n=5, respectively). F) Most abundant BMP lipid species that are significantly altered in the isolated hepatocytes with cold exposure are shown. BMP lipid abundances normalized to internal standards and tissue weight. G) Lipid abundance in whole liver across several classes are reported, as measured by untargeted lipidomics. Abundances are relative to RT controls for each lipid class. Data reported as mean±SD of experimental groups. \**p*<0.05, \*\**p*<0.01, \*\*\**p*<0.001, \*\*\*\**p*<0.0001. Abbreviations: Cers, ceramides; LPC, lysophosphatidylcholines; HexCer, hexosylceramides; BMP, bis(monoacylglycero)phosphate; Con, control; FFA, free fatty acids; KD, knockdown; RT, room temperature; TFEB, transcription factor EB; TG, triglycerides

### Pla2g15 is responsible for the cold induced changes in BMP lipids

To directly test if Pla2g15 regulates the cold induced changes in BMP lipids, we knocked down Pla2g15 in the livers of mice (Pla2g15 KD, AAV8-shPla2g15) (**Figure 6C & S6A**). Pla2g15 KD led to improved cold tolerance, with mice better able to maintain their body temperature (**Figure 6D**). To ensure that the changes we observed were driven by hepatocytes, we isolated hepatocytes and confirmed that BMP lipids were increased with loss of Pla2g15 (**Figure 6A, E, & F, S6B-C**). The improved cold tolerance with Pla2g15 KD coincided with a loss of cold induced increases in BMP lipids (**Figure 6E-F & S6D-E**). Interestingly, loss of Pla2g15 also ablated other cold induced changes in liver lipids observed including ceramides, lysophosphatidylcholine, free fatty acids, hexosylceramides, and ether lipids (**Figure 6G**).

### Pla2g15 is a BMP lipid phospholipase

To confirm that Pla2g15 is acting as a phospholipase to directly catabolize BMP lipids, we prepared liposomes containing the commercially available di-oleoyl BMP with non-hydrolyzable ether lyso-PC 18:0 in an acidic buffer to mimic the lysosome environment. We then added purified human Pla2g15 protein treated with or without the catalytic serine inhibitor diisopropyl fluorophosphate (DFP) and monitored for the degradation product oleic acid via TLC (**Figure 7A**). Fluorophosphonates like DFP block the catalytic serine residue of Pla2g15 and inhibit activity(Bouley et al., 2019; Glukhova et al., 2015). In the presence of DFP inhibitor, there was no migration band related to oleic acid, however, in the reactions of liposomes and Pla2g15 only, there was a band characteristic of oleic acid, indicating degradation of di-oleoyl BMP (**Figure 7A**).

**Figure 7:**
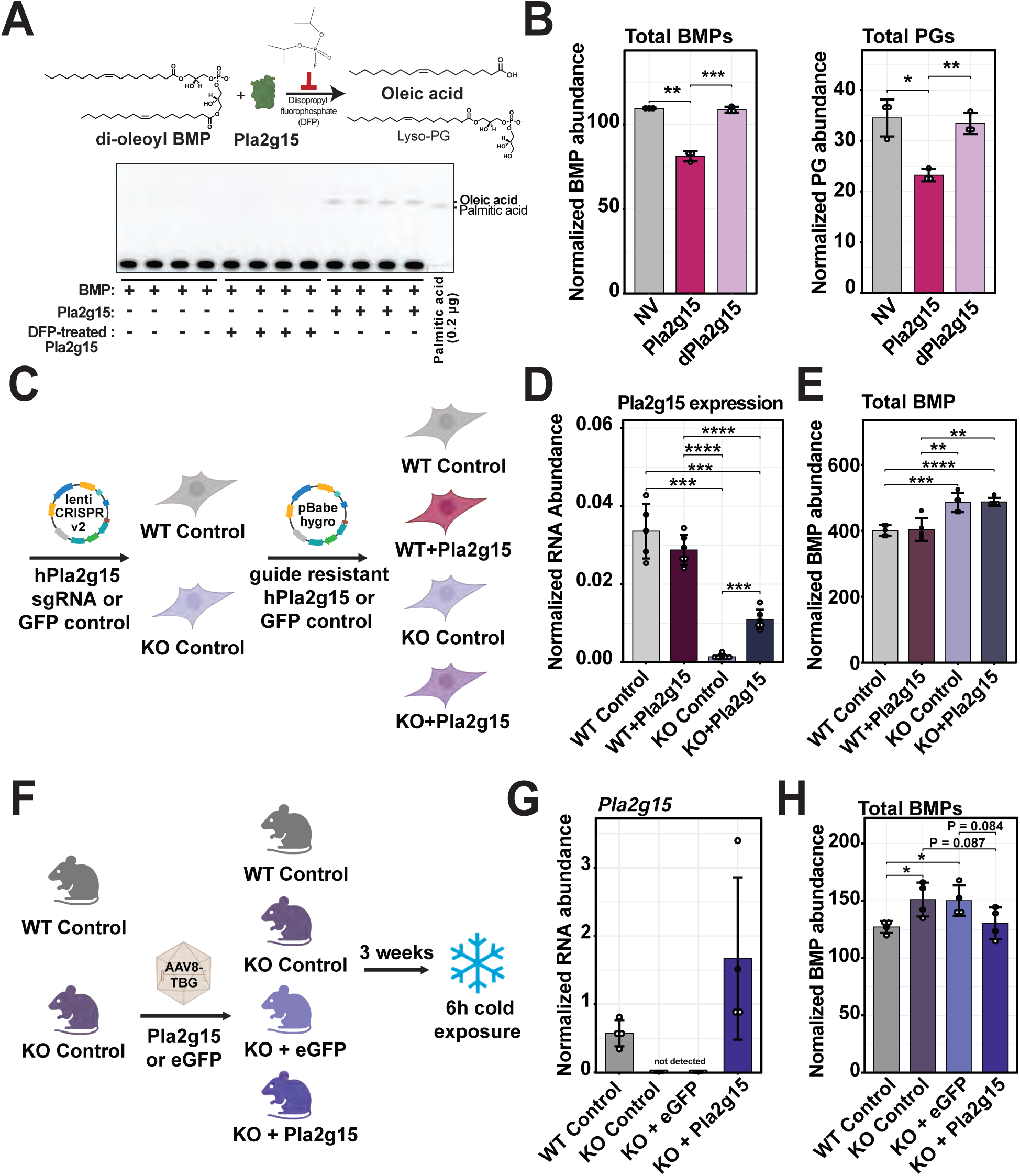
Pla2g15 degrades BMP lipids in vivo. A) Schematic of an in-vitro liposome assays using commercially available di-oleoyl BMP that was used to assess Pla2g15 activity (top). The expected products from Pla2g15 degradation of di-oleoyl BMP are oleic acid and lysophosphatidylglycerol (LPG). The Pla2g15 inhibitor diisopropyl fluorophosphate was used as negative control. In the present solvent system (see Method details), one of the products, oleic acid, migrated onto the plate and was detected after release from BMP (n=4/group). B) Total BMP lipid and PG abundance in liver lysates from mice exposed to cold for 6 hours and then incubated with bacterially expressed Pla2g15 or catalytically dead Pla2g15 (dPla2g15; n=3/group). Bacteria expressing no vector (NV) was used as a control. C) Experimental design for generating Pla2g15 knockout (KO) 293T cells and overexpressing Pla2g15 in the KO background. D) Pla2g15 gene expression is shown for the Pla2g15 KO, re-expression, and control cell lines described in C (n=5/group). E) Total BMP lipid abundance measured in 293T cells with Pla2g15 KO and overexpression. F) Schematic of experimental design for overexpressing Pla2g15 via AAV8-TBG in wild type (WT) and mice lacking Pla2g15 (KO). Three weeks post AAV injection, mice were kept at cold (4°C) for 6 hours (n=4/group). G) Hepatic expression of Pla2g15 are plotted for the mice described in F, after exposure to 6h of cold. H) Total BMP lipids are reported, as measured by targeted lipidomics in the livers of control and Pla2g15 KO mice treated with an eGFP control or Pla2g15 AAV and exposed to cold for 6 hours. Lipids are normalized to internal standards and tissue weight. When relevant, mean±SD is plotted. \**p*<0.05, \*\**p*<0.01, \*\*\**p*<0.001, \*\*\*\**p*<0.0001. Abbreviations: DFP, Diisopropyl fluorophosphate; PG, phosphatidylglycerol; NV, no vector; dPla2g15, catalytically dead Pla2g15; WT, wild-type; KO, knockout.

Phospholipase A2 enzymes contain a catalytic triad of a serine, histidine, and aspartic acid that facilitates the hydrolysis of the ester bond typically in the sn-2 position(Burke and Dennis, 2009; Shayman and Tesmer, 2019). To determine if Pla2g15 was directly acting on BMP lipids through this catalytic site, we mutated serine 165 to an alanine to generate a catalytically dead Pla2g15 (dPla2g15). Wild-type murine Pla2g15 and dPla2g15 were expressed in bacteria, isolated, and then incubated in liver lysate from cold exposed mice for 15 minutes. Pla2g15 incubation led to a decrease in endogenous BMP lipids and PG, while dPla2g15 did not alter BMP lipid levels (**Figure 7B & S7A**). We also generated Pla2g15 knockout in 293T cells and rescued expression with a Cas9 resistant Pla2g15— although expression was only partially restored (**Figure 7C & D**). Loss of Pla2g15 increased total BMP lipid species (**Figure 7E**), and rescue with Cas9 resistant Pla2g15 decreased several BMP lipids including 18:1_18:1 and 18:0_18:1 (**SFigure 7B**).

To determine if Pla2g15 KO directly altered liver BMP lipid levels, we assessed control (*Pla2g15+/+*) and knockout (*Pla2g15-/-*) mice(Akira et al., 2007; Hiraoka et al., 2006). Because this is a whole body knockout and there can be compensation, we re-expressed Pla2g15 in the hepatocyte alone with hepatocyte specific driver thyroxine binding globulin (AAV8-TBG Pla2g15) three weeks before cold exposure (**Figure 7F&G**). Mice were placed in cold for 6 hours to increase BMP lipids. Pla2g15 KO mice had elevated BMP lipid levels relative to their litter mate controls, and re-expression of Pla2g15 in hepatocytes decreased BMP lipid levels (**Figure 7G & H, S7C**).

## Discussion

Response to cold exposure is metabolically challenging and requires coordination of metabolism across multiple organs. The liver plays a central role in the thermogenic response, altering lipid metabolism to fuel heat production(Bornstein et al., 2023; Simcox et al., 2017; Zhang et al., 2024). Through lipidomics on the liver, we were able to determine that BMP lipids are increased during cold exposure. Our studies determined that the rise in BMP lipids is driven by inhibition of Pla2g15, a BMP lipase that is transcriptionally controlled by TFEB. During cold exposure TFEB has altered DNA occupancy, predominantly regulating genes associated with lipid metabolism. We also observed a decrease of BMP lipids in the liver of TFEB KD mice exposed to cold coinciding with cold intolerance. Together, these studies provide new insights into liver thermoregulatory processes and the regulation of BMP lipids. BMP lipid levels are altered in a number of human diseases including hepatic steatosis and lysosomal storage disorders, although the regulation of BMP lipids in these diseases is poorly understood(Akira et al., 2007; Shayman and Tesmer, 2019).

There has been a resurgence in BMP lipid research since their initial characterization in 1967. This resurgence has been driven by improved methods to detect BMP lipids(Anderson et al., 2017; Body and Gray, 1967; Scherer et al., 2010), the potential for BMP lipids as biomarkers of metabolic and neurodegenerative disease(Boland et al., 2022; Grabner et al., 2019; Grabner et al., 2020; Logan et al., 2021), and the understanding that BMP lipid regulate lysosomal function. In the past two years, CLN5 and PLD3/4 (Bulfon et al., 2024; Medoh et al., 2023; Singh et al., 2024) have been established as regulators of BMP lipid synthesis and are important in esterification of lysophosphatidylglycerol to BMPs. Our work adds to this growing exploration by functionally characterizing Pla2g15 as an enzyme capable of degrading BMP lipids, and establishes regulation of Pla2g15 by TFEB. Understanding mechanisms to regulate BMP lipid levels can yield therapeutic potential since treatment of NPC1 deficient cells with BMP lipids alleviates cholesterol burden and reduces cellular toxicity, which are characteristic of NPC diseases(Alvarez-Valadez et al., 2024; Ilnytska et al., 2021).

Pla2g15 was originally identified as a lyso-phosphatidylcholine lipase and a transacylase that catalyzed the formation of 1-O-acylceramides from sn-2 fatty acyl groups(Abe et al., 2006, 2007; Taniyama et al., 1999). In the absence of ceramide acceptors, Pla2g15 exhibits phospholipase activity against a wide range of glycerophospholipids and oxidized phospholipids(Abe et al., 2006; Hinkovska-Galcheva et al., 2018; Nyame et al., 2024a). The enzyme contains a core α/β-hydrolytic domain and catalytic triad consisting of Ser165, Asp327, and His359. The potential for Pla2g15 to regulate BMP lipids was first established by a recent study that identified it as a putative phosphatidylglycerol deacylase that contributes to BMP synthesis in HeLa cells(Chen et al., 2023). More recent work identified and enzymatically characterized Pla2g15 as a BMP phospholipase using BMP lipid standards which we were able to recapitulate (**Figure 7A**) (Abe et al., 2024). Our work and the work of the Abu-Remaileh lab utilized assays in liver lysate to demonstrate that Pla2g15 is a BMP phospholipase. We confirmed these results in multiple cell models, Pla2g15 knockout mice, and in Pla2g15 knockdown mice(Nyame et al., 2024b). While our studies indicate a potential function of elevated BMP lipids in energy homeostasis in Pla2g15 KD mice having improved cold tolerance, Abu-Remaileh lab’s further went on to characterize the importance of Pla2g15 in lysosomal storage disorders.

There are over 300 single nucleotide variants that have been characterized for *Pla2g15(Chen et al., 2024)*; additional work to understand Pla2g15 function and regulation will be crucial for the exploration of metabolic health and lysosomal storage disorders. Evidence is emerging that Pla2g15 may be important in autophagy(Kagohashi et al., 2023; Li et al., 2022), lysosomal storage disorders(Nyame et al., 2024a; Nyame et al., 2024b), and energy homeostasis. Further exploration of the mechanism by which TFEB activity is altered to regulate Pla2g15 and the development of agonists for Pla2g15 are needed to probe the therapeutic potential of these discoveries.

## Acknowledgments

We would like to thank Edrees H. Rashan, CiJi Lim, and Alan Attie for advice on experimental design and all members of the Simcox Lab for assistance with mouse experiments, editing materials, and thoughtful discussions. We also thank the Lim Lab at the UW-Madison for equipment use. Dr. Abu-Remaileh is a Stanford Terman Fellow and a Pew-Stewart Scholar for Cancer Research, supported by the Pew Charitable Trusts and the Alexander and Margaret Stewart Trust. Kwamina Nyame is supported by the Sarafan ChEM-H Chemistry/Biology Interface Program as Kolluri Fellow. Kwamina is additionally supported by the Bio-X Stanford Interdisciplinary Graduate Fellowship affiliated with the Wu Tsai Neurosciences Institute (Bio-X SIGF: Mark and Mary Steven’s Interdisciplinary Graduate Fellow). Dr. Simcox is an HHMI Freeman Hrabowski Scholar. Research reported in this publication was supported by the Glenn Foundation and American Federation for Aging Research (A22068 to JS); Hatch Grant (WIS04000-1024796 to JS and RJ); JDRF (JDRF201309442 to JS); an R01 through NIH/NIDDK (R01DK133479 to JS); the Dr. Stephen Babcock AG Chem Fellowship (IJ); NIH (P41 GM108538 to JJC). This material is based upon work supported by the National Science Foundation Graduate Research Fellowship Program under Grant Nos. DGE-2137424 to JWD and DGE-1747503 to HV as well as support of the NIH/NIGMS funded Biotechnology Program T32 (5T32GM135066-05;TK & AH). Any opinions, findings, and conclusions or recommendations expressed in this material are those of the author(s) and do not necessarily reflect the views of the National Science Foundation. The content is solely the responsibility of the authors and does not necessarily represent the official views of the National Institutes of Health. The work was also supported in part by startup funds from the University of Wisconsin-Madison School Department of Biochemistry to JS.

## Author Contributions

Conception: JD, RJ, and JAS; methodology: JD, RJ, TK, GG, DJN, AA, AC,KN, AH, SM, XC, RV, IJ, KS, VHG, HV, DAB, GW, and ES; original draft: RJ and JAS; revisions: JD, TK, RJ, AC, IJ, and JAS; materials: WXD, JJC, JS, MA, and JAS; supervision: JS, MA, JAS.

## Supplementary Figures

Figure S1. BMP lipids are altered during acute cold exposure in the liver and in circulation in mice. A) Chromatograph traces of untargeted, LC/MS-based lipidomics is shown for liver. Identified lipid classes are labeled above the chromatographic regions of elution, with lipids predominantly identified in positive ionization in orange, and negative ionization in brown. B) Chromatographic separation and fragmentation spectra and structures of isomeric BMP and PG species containing 18:1/18:1 acyl chains are shown. Standards were purchased from Avanti Polar Lipids. A targeted LC/MS method was developed to distinguish and quantify BMP and PG species. C) BMP lipid species unchanged between RT (fasted for 6h) and cold exposed (4°C for 6h) male mice (n=5-6/group) are shown. D and E) Total and individual PG species from liver measurements of the same mice as in Fig S1B are shown. F) Total and species-specific changes in plasma levels of BMP and PG lipids for mice from Fig S1B are plotted. G) Total BMP and PG lipid changes in the liver of mice kept at RT (fasted) or cold exposed (4°C) over a 6 hour (n=3/group) and a 24 hour (n=5/group) timecourse. H) Total BMP and PG lipid changes in the liver of mice housed at either RT or cold (4°C) for 1 week and given ad libitum access to food and water (n=6/group). Lipid abundances are normalized to internal standards and tissue weight or plasma volume. \**p*<0.05, \*\**p*<0.01, \*\*\**p*<0.001. Abbreviations: BMP, bis(monoacylglycero)phosphate; LC/MS, liquid chromatography mass spectrometry; PG, phosphatidylglycerol; RT, room temperature.

Figure S2. Liver transcripts of lysosome-associated processes are altered during acute cold. A - F) Data for all transcripts measured in Fig 3A are plotted. Mice were fasted and kept at either RT or cold (4°C) for 6h. Gene expression was measured for 6 mice/group and only transcripts with CT values < 30 were reported. G) BMP lipid measurements on lysosomal IP from the livers of LysoTag mice exposed to RT of cold (4°C) for 6h (n=4-5/group). \**p*<0.05, \*\**p*<0.01, \*\*\**p*<0.001. Abbreviations: RT, room temperature.

Figure S3. TFEB liver knockdown caused lysosomal alterations and changes in circulating fuel, leading to cold intolerance. A) Total liver BMP lipids were measured from male B6 mice treated with an AAV8 harboring shRNA to *Tfeb* or scramble sequence as control. Mice were housed at RT and with *ad libitum* access to food and water prior to measurements (n=5/group). B) Body weight was measured before and after mice treated with control or an AAV8 virus with shRNA targeting Tfeb were housed at RT or cold (4°C) for 6h (n=6/group). Change in body weight was compared (left), as well as wet liver weight (right) at the end of the experiment. C) Hepatic and circulating levels of acylcarnitines were measured in mice from Figure S3B. D) *Tfeb* gene expression was measured in various tissues in mice from Figure S3B. E) Gene expression of liver transcripts related to mitochondrial and lysosomal abundance and activity were measured in mice. Technical replicates were assessed for all transcripts. F) Protein expression of lysosome and autophagy-related proteins was assessed in the liver of mice from Fig S3B using western blot. Densitometry results are plotted. G) Circulating ketones (left) and fatty acids in the liver (right) were measured in mice with or without TFEB knockdown (KD) that were kept at either RT or in the cold for 6 hours (n=6/group). \**p*<0.05, \*\**p*<0.01, \*\*\**p*<0.001. Abbreviations: AAV8, adeno-associated virus serotype 8; ACars, total acylcarnitines; BHB, β-hydroxybutyrate; NEFA, non-esterified fatty acid; RT, room temperature; TFEB, transcription factor EB.

Figure S4. ChIP-seq of liver TFEB indicates a unique program during cold exposure. A) Histograms comparing the distance from transcription start sites (TSS) of ChIP-seq analysis of liver TFEB from RT and 6h fasted, cold-exposed (4°C) mice was compared (*q*<0.05 for all sites used in comparison). B) The type of gene associated to TFEB binding sites in RT (top) or cold (bottom) were compared. C) Targeted ChIP-qPCR was performed on lipid metabolism-associated genes with TFEB binding site from ChIP-seq analysis was used to validate findings. TFEB ChIP-qPCR from liver (n=2) was compared to IgG (n=2) for validation. Relative enrichment of target binding sites is shown on the left with the associated ChIP tracks on the right. An arrow denotes the binding region for which qPCR primers were designed. D) Proteomic analysis of interactors that co-immunoprecipitate with TFEB in the liver of mice exposed to either RT or cold for 6h (n=3). Annotations reported were present in all 3 samples. \**p*<0.05, \*\**p*<0.01, \*\*\**p*<0.001, \*\*\*\**p*<0.0001. Abbreviations: ChIP, chromatin immunoprecipitation and sequencing; RT, room temperature; TFEB, transcription factor EB.

Figure S5. BMP lipids are regulated by the lysosomal trafficking and gene expression. A) Gene expression of human *Tfeb* (h*Tfeb,* left) and total BMP lipids (right) in Hepa1-6 cells with retroviral overexpression of *hTfeb* or *LacZ* as control (n=6/group).Total BMPs were normalized to protein amount from cells. B) Gene expression of mouse *M6pr* (left) in Hepa1-6 cells treated with siRNA containing a scramble sequence or targeting murine *M6pr* (n=4/group; technical replicates), which traffics lysosomal proteins including Pla2g15 to the lysosomes. Total BMP lipids (right) were measured in the Hepa1-6 cells treated with siRNA with scramble sequence (n=3) or murine *M6pr* (n=5). Lipid data was normalized to internal standards and total protein. \**p*<0.05, \*\**p*<0.01, \*\*\**p*<0.001, \*\*\*\**p*<0.0001. Abbreviations: BMP, bis(monoacylglycero)phosphate; KD, knockdown; *M6pr*, mannose-6-phosphate receptor; RT, room temperature; TFEB, transcription factor EB.

Figure S6. *Pla2g15* is a regulator of BMP lipids in the liver. A) Gene expression for *Pla2g15* in whole liver from mice three weeks post treatment with AAV8-GFP-U6-scrmb-shRNA (control; n=12) or AAV8-GFP-U6-mPla2g15-shRNA (Pla2g15 knockdown (KD); n=12) and immediately following 6h at either RT (n=6/group) or cold (n=6/group). B) Gene expression for albumin and CPT1a in isolated hepatocytes from mice described in figure 6C, treated with control or AAV8 containing shRNA targeting Pla2g15 and subjected to 6h at RT or cold. C) Select BMP lipid species were measured in isolated hepatocytes described in figure 6C. D) Total BMP and PG lipid abundances from mice described in Figure S6 are plotted (n=6/group). E) Lower abundance BMP lipid species were measured in whole livers of the mice described in Figure S6. All lipid abundances were normalized to internal standards and either tissue weight (whole liver) or protein content (isolated hepatocytes). \**p*<0.05, \*\**p*<0.01, \*\*\**p*<0.001, \*\*\*\**p*<0.0001. Abbreviations: RT, room temperature; KD, knockdown.

Figure S7. Pla2g15 is a degrader of BMP lipids. A) BMP lipid species abundances in the liver lysate incubated with bacterially expressed Pla2g15 or catalytically dead Pla2g15 (dPLa2g15), described in Figure 7B (n=3/group). Bacteria expressing no vector (NV) was used as a control. Lipid abundances were normalized to internal standards and protein content. B) Abundances of BMP lipids in WT and Pla2g15 knockout (KO) 293T cells with overexpression of Pla2g15 (+ Pla2g15) are plotted, as described in Figure 7C (n=5/group). C) Select BMP lipid species measured in the liver of WT, Pla2g15 KO mice, and Pla2g15 KO mice treated with an AAV8-TBG-eGFP or AAV8-TBG-Pla2g15. The experimental design is provided in Figure 7F (n=4/group). D) Total PGs in the liver of the mice described in Figure 7F are reported. BMP and PG lipid abundances are normalized to internal standards and tissue weight. \**p*<0.05, \*\**p*<0.01, \*\*\**p*<0.001, \*\*\*\**p*<0.0001. Abbreviations: NV, no vector; dPla2g15, catalytically dead Pla2g15; WT, wild-type; KO, knockout.

## Supplementary Tables

Tables S1. Sequences for all qPCR primers.

## STAR Methods

| REAGENT or RESOURCE | SOURCE | IDENTIFIER |
| --- | --- | --- |
| **Antibodies** |  |  |
| Rabbit anti-Pla2g15 | Novus Biologicals | NBP2-17193 |
| Rabbit anti-TFEB (western blot) | Bethyl Laboratories | A303-673A |
| Tfeb polyclonal antibody | Proteintech | 13372-1-AP |
| TFEB rabbit mAb (IP) | Cell Signaling | 83010 |
| LC3B Antibody | Cell Signaling | 2775 |
| Normal Rabbit IgG | Cell Signaling | 2729 |
| Phospho-(Ser/Thr) Phe Antibody | Cell Signaling | 9631 |
| Histone H3 (1B1B2) Mouse mAb | Cell Signaling | 14269 |
| Rab5a polyclonal antibody | Proteintech | 11947-1-AP |
| LIPA polyclonal antibody | Proteintech | 12956-1-AP |
| CD107B/LAMP2 monoclonal antibody | Proteintech | 66301-1-Ig |
| Human/mouse/rat beta-actin antibody | R&D Systems | MAB8929 |
| Recombinant anti-HNF-4-alpha antibody | Abcam | ab181604 |
| Anti-LAMP1 | Development Studies Hybridoma Bank, Iowa | 1D4B |
| Goat anti-Rabbit IgG Secondary antibody HRP | Fisher Scientific | PI32460 |
| Goat anti-Mouse IgG Secondary antibody HRP | ThermoFisher Scientific | A-10668 |
| Goat anti-Rat IgG Secondary antibody HRP | Fisher Scientific | A10549 |
| **Chemicals, peptides, and recombinant proteins** |  |  |
| Splash II Lipidomix | Avanti Polar Lipids | 330709 |
| 14:0 BMP (S, R) | Avanti Polar Lipids | 857131 |
| 18:1 BMP (S, R) | Avanti Polar Lipids | 857133 |
| 18:1 (delta-9-cis) PG | Avanti Polar Lipids | 840475 |
| 15:0-18:1-d7-PG | Avanti Polar Lipids | 791640 |
| 15:0-18:1-d7-PA | Avanti Polar Lipids | 791642 |
| C15 Ceramide-d7 (d18:1-d7/15:0) | Avanti Polar Lipids | 860681 |
| Stearic acid-d35 | Cayman Chemical | 9003318 |
| Oleoyl-L-carnitine-d3 (chloride) | Cayman Chemical | 26578 |
| Pierce Formic Acid, LC-MS Grade | ThermoFisher Scientific | 85178 |
| Acetonitrile | Honeywell | LC015 |
| Ammonium Formate | Honeywell | 55674 |
| Water B&J Brand, LC-MS | Fisher Scientific | 600-30-78 |
| Fisher Chemical Isopropanol Optima, LC-MS | Fisher Scientific | A4614 |
| DMEM | ThermoFisher Scientific | 11-965-118 |
| Fetal Bovine Serum (FBS) | ThermoFisher Scientific | 10-437-028 |
| DPBS | ThermoFisher Scientific | 14190144 |
| Lipofectamine RNAiMAX | ThermoFisher Scientific | 13778030 |
| Opti-MEM | ThermoFisher Scientific | 31985062 |
| Laemmli sample buffer | Bio-Rad | 161-0737 |
| β-mercaptoethanol | Sigma-Aldrich | M6250 |
| RIPA lysis buffer | Fisher-Scientific | NC984499 |
| Pierce protease inhibitor mini tablets, EDTA free | Fisher-Scientific | PIA32955 |
| 4-15% Mini-Protean TGX precast protein gels | Bio-Rad | 4561096 |
| PowerUp SYBR Green Master Mix | ThermoFisher Scientific | A25778 |
| TRIzol | Invitrogen | 15596018 |
| Cathepsin B substrate III – Fluorogenic, Calbiochem | Millipore Sigma | 219392 |
| E-64-D | Enzo Life Sciences | BML-PI107-0001 |
| Isoflurane 99.9% | McKesson | 803250 |
| 25% glutaraldehyde | Electron Microscopy Sciences | 16220 |
| 0.4M Sodium cacodylate | Electron Microscopy Sciences | 11655 |
| **Critical commercial assays** |  |  |
| Glucosylceramidase activity assay kit | Abcam | ab273339 |
| β-Hydroxybutyrate colorimetric assay kit | Cayman Chemical | 700190 |
| Wako HR Series NEFA kit | Fujifilm | 999-34691; 995-34791; 991-34891; 993-35191; 276-76491; 997-76491 |
| SimpleChIP Plus kit (Magnetic Beads) 4°C and RT reagents | Cell Signaling | #38191 |
| Pierce Protein A/G Magnetic Beads | VWR | PI88802 |
| Pierce BCA Protein Assay | ThermoFisher Scientific | #23225 |
| High Capacity cDNA Reverse Transcription Kit | ThermoFisher Scientific | #4368813 |
| Zymo ChIP DNA Clean and Concentrator Kit | VWR | D5205 |
| Qubit dsDNA HS Assay Kit | ThermoFisher Scientific | Q32851 |
| Agilent DNA HS Kit | Agilent Technologies | 5067-4626 |
| TruSeq ChIP Sample Preparation kit | Illumina | IP-202-1012 |
| SuperSignal West Atto Ultimate Sensitivity Substrate | ThermoFisher Scientific | A38554 |
| **Deposited data** |  |  |
| Untargeted lipidomics data (WT mice) | https://doi.org/10.1016/j.jlr.2022.100197 | MTBLS3730 |
| Targeted LC/MS lipidomics data | This paper | ftp://massive.ucsd.edu/v06/MSV000096791/ |
| TFEB ChIP-seq | This paper | GEO |
| **Experimental models: Cell lines** |  |  |
| Hepa1-6 cells | ATCC | CRL-1830 |
| **Experimental models: Organisms/strains** |  |  |
| Mouse: C57BL/6J male mice 9 weeks | Jackson Laboratories | #000664 |
| **Oligonucleotides** |  |  |
| ON-TARGETplus Mouse Tfeb siRNA | Dharmacon | L-050607-02-0005 |
| ON-TARGETplus Non-targeting Control Pool | Dharmacon | D-001810-10-05 |
| ON-TARGETplus Mouse M6pr siRNA | Dharmacon | L-050312-01-0005 |
| ON-TARGETplus Mouse Pla2g15 siRNA | Dharmacon | L-059873-01-0005 |
| Primers see Supplementary Table 1 |  |  |
| **Recombinant DNA** |  |  |
| AAV8-GFP-U6-scrmb-shRNA | Vector Biolabs | 220207-220729 |
| AAV8-GFP-U6-mTFEB-shRNA | Vector Biolabs | 220801#38 |
| **Software and algorithms** |  |  |
| R version 4.3.0 | R Core team | https://www.r-project.org |
| Agilent MassHunter Suite | Agilent Technologies | G3337AA; M5960AA; M6010AA |
| Agilent LipidAnnotator | Agilent Technologies | G3835AA |
| LipidCreator | Peng *et al.* | https://doi.org/10.1038/s41467-020-15960-z |
| R scripts for data analysis | This paper | Github: RJain52 |
| **Other** |  |  |
| TissueLyser II | Qiagen | 85300 |
| Tecan Infinite M Plex | Tecan | 30190085 |
| QuantStudio 5 Real-Time PCR System | ThermoFisher Scientific | A34322 |
| Acquity UPLC BEH C18 Column | Waters Corporation | 186002352 |
| Acquity UPLC BEH C18 VanGuard Pre-Column | Waters Corporation | 1860003975 |
| Agilent 1290 Infinity II LC System | Agilent Technologies | - |
| Agilent 6495c triple quadrupole mass spectrometer | Agilent Technologies | G6495CA |
| Agilent 6546 Q-TOF mass spectrometer | Agilent Technologies | G6546AA |
| iBright CL1500 Imaging System | ThermoFisher Scientific | A44240 |
| Amersham Protran nitrocellulose blotting membrane | Fisher | 45004003 |
| Physitemp rectal probe for mouse | Fisher | NC9713069 |
| JEM 1016CX electron microscope | JEOL Ltd. | - |
| Formulab Diet | LabDiet | 5008 |

## Method Details

### Mouse husbandry

All animal experiments were performed in accordance with institutional guidelines and were approved by the Institutional Animal Care and Use Committee at the University of Wisconsin-Madison. Mice were housed at room temperature (21-24°F), with relative humitidy ranging from 30-50% with an average of 37%, using 12hr dark and 12hr light cycles. Mice had *ad libitum* access to water and standard chow (Rodent Diet 5001) prior to experiments. Age, sex, and treatment are specified in each figure legend. *Pla2g15* control *(Pla2g15+/+)* and knockout *(Pla2g15-/-)* mice were maintained as litter mate controls through breeding heterozygous mice (*Pla2g15+/-)*. Genotyping was performed at 21 days, on tail clipping, by Transnetyx using previously published methods and primers, 5′-CAGGGTAGCTCACAACTCTTTG-3′; b, 5′-CAAAGCTCTGGACTGTTTTCCTGC-3′; c, 5’-GAATTCCTAGACCCCAGCAAGAGGAATGTG-3′; and d, 5′-CCCTCCCCAGAGATGGATATTT-3′. (Akira et al., 2007; Hiraoka et al., 2006). LysoTag mice were maintained in accordance with the Institutional Animal Care and Use Committee at Stanford University(Laqtom et al., 2022).

### Cold tolerance test

Mice were placed without bedding at room temperature (RT; 22°C) or cold (4°C), food was removed at the start of the experiment, and mice had *ad libitum* access to water (n=5-6/group). Temperature was taken hourly via rectal probe and mice were monitored for any changes in behavior or physiology. Experiments started at zeitgeber time=3. At the end of the CTT, isoflurane as administered and mice were euthanized post-anesthesia via cervical dislocation. Plasma was obtained via cardiac puncture; post-euthanasia tissue was harvested, washed with PBS, sectioned and either flash frozen in liquid nitrogen or further processed. Hepatocyte isolations were performed immediately following the CTT following standard procedure (Severgnini et al., 2012).

### AAV injection

For WT, *Tfeb* KD, *Pla2g15* KD, and *Pla2g15* KO experiments, littermate controls were used and mice were age and weight-matched. Mice were retro-orbitally injected with the appropriate AAV at 1e11 genome copies/mouse (mice weighing 23-28g). Post-injection weights were monitored to ensure no deviation between groups and that no more than 20% body weight was lost post-injection. Subsequent experiments were performed three week after injections.

### Cell culture

Hepa1-6 cells were cultured at 5% CO2 at 37°C. Cells were maintained in DMEM with 4.5 g/L glucose, supplemented with 10% fetal bovine serum. For knockdown of *Tfeb* and *Pla2g15*, ON-TARGETplus Mouse Tfeb pooled siRNA was purchased from Horizon Discovery. Transfection was performed in Hepa1-6 cells in a 6-well plate using 50 pmol siRNA and Lipofectamine RNAiMAX according to the manufacturer’s instructions. Briefly, 5e5 cells were plated per well in a 6-well culture plate. Cells were cultured in DMEM+10% FBS overnight. The next day, media was replenished and a mixture of 300μL Opti-MEM with 9uL lipofectamine agent and appropriate siRNA was added to wells. Media was replenished 24h later to remove siRNA, then harvested after an additional 24h of culturing.

### Generation of Pla2g15 knockout and overexpression cell lines

Knockouts were generated in Hek 293T cells using a CRISPR-Cas9 lentiviral system. Briefly 5 x 106 cells were transfected with the helper plasmids pVSVg, pMDLgRRE, pRSV-rev and the pLenti plasmids containing either the Pla2g15 guide or a GFP control. Transfection was carried out using the Lipofectamine 3000 reagent (Thermo Scientific #L3000008). After 2 days virus was harvested, filtered through a 0.45um syringe filter and 1uL of 10mg/mL polybrene was added per mL of media harvested. Virus was added to Hek 293T cells and selected for puromycin resistance (3uL of 10mg/mL puro per 10mL of media) after 3 days. For Pla2g15 overexpression in the knockout cell lines, PhoA cells were grown to 80% confluence and transfected with pBABE plasmids and PEI transfection reagent. After 2 days virus was harvested, filtered through a 0.45um syringe filter and 10mg/mL polybrene was added at 1:1000. Virus was added to 293T cells and selected for hygromycin resistance (200ug/mL).

### Luciferase Assay

Potential TFEB binding sites were identified from ChIP data and aligned with mm10 reference genome (chr8: 106143980-106151020). This sequence was divided into 5 regions of ∼1500bp, with ∼100bp overlap and each region was cloned into a pGL3 Basic luciferase reporter plasmid (pGL3 Pla2g15 R1-R5). 293T cells overexpressing LacZ or TFEB were generated using a lentiviral system (pLenti-CMV). These 293T cells were plated in 96-well plates and transfected with with pGL3 Basic empty vector, pPparg (addgene#8896), or pGL3 Pla2g15 R1-R5, and pCMV-Renilla control plasmid (addgene#118066). After 48h of growth at 37°C, Dual-Glo Luciferase Assay System (Promega, E2940) was used to assess luciferase and Renilla activity. Final activity was expressed as Luciferase Activity/Renilla Activity.

### RNA extraction and RT-PCR

RNA was recovered from tissues or cells with TRIzol reagent followed by homogenization using TissueLyser. cDNA was synthesized using High-Capacity cDNA Reverse Transcription Kit. RT-PCR amplification was for 40 cycles using PowerUp SYBR Green Master Mix in a 384-well plate using QuantStudio 5 Real-Time PCR System. Primer sequences are listed in **Table S1**. For quantification, a standard curve was generated for each primer set and used to determine relative gene expression. Expression levels are presented relative to the housekeeping gene *Rps3*.

### ChIP-seq

Chromatin immunoprecipitation was performed on livers from individually housed male C57BL/6J mice (11 weeks old) that were placed at 4°C without food for 6 hours or kept at RT with ad libitum access to food (n=2 per group). All mice had ad libitum access to water. SimpleChIP Plus Enzymatic Chromatin IP Kit was used according to manufacturer’s instructions. Briefly, 100mg of tissue was used per mouse and two immunoprecipitations were performed with rabbit anti-TFEB antibody, and two immunoprecipitations were performed with normal rabbit IgG control. ChIP DNA was purified using Zymo ChIP DNA Clean and Concentrator Kit. Purified immunoprecipitated and input DNA was quantified and assessed for sheering using the Qubit dsDNA HS Assay Kit and Agilent DNA HS chip, respectively. Samples were prepared according the TruSeq ChIP Sample Preparation kit. Libraries were size selected for an average insert size of 350 bp using SPRI-based bead selection. Quality and quantity of the finished libraries were assessed using an Agilent Tapestation and Qubit dsDNA HS Assay Kit, respectively. Paired end 150bp sequencing was performed on the NovaSeq 6000 by the UW-Madison Biotechnology Center.

Sequenced reads were checked for: sequencing quality, insert length, read concordance, duplications, and adapters via FastP and then aligned to the mouse genome (UCSC build mm10) via bowtie2 v2.2.5 using the sensitive end-to-end read alignment mode. Aligned reads are sorted and indexed with SAMtools 1.9 and peaks are called at *p*≤0.05 and *q*≤0.05 (separately) using the input/IgG as the background control with the MACS2. Primary and differential peaks were annotated for their positioning relative to genomic elements with Homer v4.11. *De novo* and known motif analysis were carried out with Homer. Pathway ontology information was generated by inputting genes differentially bound in cold or RT into geneontology.org for ‘Reactome’ pathway results.

### TFEB Immunoprecipitation

Liver (100mg) was homogenized in RIPA buffer with protease and phosphatase inhibitors using the TissueLyzer II in round-bottom microcentrifuge tubes with a single steel bead. Homogenate was centrifuged at 4°C for 15min at 16000 RPM and the supernatant was transferred to a new tube. This process was repeated to pellet additional cell debris. Protein concentration was measured by BCA assay, and 1mg of protein was aliquoted into low protein-binding tubes with 1uL anti-TFEB antibody (Proteintech 13372-1), 5uL anti histone H3, or anti IgG. Volume was adjusted to 500uL with RIPA buffer, and samples were mixed overnight at 4°C.

Protein A/G magnetic beads were then added to the samples, which were allowed to mix for 4 hours at 4°C. Beads were washed in RIPA buffer. For Western blot, beads were resuspended in Laemmli sample buffer with 5% β-mercaptoethanol and heated at 95°C for 5min. Beads were removed, and the sample was transferred to low protein-binding tubes and stored at -20°C until use. For proteomics, beads were washed again in 20mM Tris 100mM NaCl pH 8, and then resuspended in 100uL 20mM Tris 100mM NaCl pH 8 and stored at -80°C until use.

### Western blot

Tissue (25-50mg) was homogenized in a round bottom microfuge tubes containing a single steel bead in RIPA buffer with protease inhibitor using the TissueLyzer II. Lysate was centrifuged at 4°C for 10min at 14000 RPM and the supernatant was transferred to a new tube. BCA assay was used to determine protein concentration, then samples were prepared at a concentration of 2mg/mL in Laemmli sample buffer containing 5% β-mercaptoethanol. Samples were heated at 95°C for 5min, then 15-30ug protein were loaded into precast gels. Gels were run at 150V for 30-45min, then transferred at 100V for 75min onto nitrocellulose membranes.

Membranes were blocked in either 5% milk or 5% BSA, per manufacturer instructions, prior to overnight incubation in the appropriate primary antibody at 1:1000 dilution in blocking buffer, overnight at 4°C. Secondary antibodies were diluted at 1:5000 in blocking buffer and incubation was performed for 1h at RT, then membranes were developed using ECL reagent. Blots were imaged and densitometry was performed on an iBright CL1500 imaging system.

### Electron microscopy

Mice were sacrificed using isoflurane and cervical dislocation, the liver was immediately perfused with 30mL PBS, followed by the fixation buffer containing 2.5% glutaraldehyde in 0.4M sodium cacodylate (pH=7.4). The liver was excised, submerged in fixation buffer, and cut into small pieces before being placed in fresh fixation buffer overnight at 4°C. The following day, samples were transferred to 15mL tube containing fresh fixation buffer and further processed for imaging. Images were collected on a JEM 1016CX electron microscope55.

### Lipid extraction

Murine tissue were extracted for lipids using a one-phase method as previously described6. For liver, 25-30mg tissue were homogenized in ceramic bead tubes in a 3:1:6 IPA:H2O:EtOAc^-^ solution containing the following internal standards (IS): 12uL of SPLASH II LipidoMix and 10μl of 10uM PG 15:0/18:1d7, 10uM BMP 14:0_14:0, 30uM Cer d18:1_15:0d7, 30uM ACar 18:1d3, 225uM C18:0d35, and 1.1mM PA 15:0/18:1d7. A TissueLyser II was used for bead beating. Homogenates were incubated at -20°C for 10min, then centrifuged at 16,000xg for 10min at 4°C to precipitate protein before the lipid containing solvent layer was transferred to a new tube. Samples were respun as previously to precipitate remaining debris, transferred to another tube, then dried down in a SpeedVac. Lipid extracts were resuspended in 150μl 100% MeOH for negative or 9:1 MeOH:toluene for positive ionization analysis. Plasma samples were extracted identically expect 50μl plasma were used as starting material. Samples were transferred for autosampler vials containing glass inserts for analysis.

Samples from cell culture were prepared by first aspirating media, washing twice with DPBS on ice, then gently scraping samples into 1mL DPBS and transferring to a microfuge tube. After pelleting the samples and aspirating DPBS, 100% MeOH containing IS as above was added to samples for lipid extraction. Cell pellets were dispersed with light vortexing and incubated on ice for 1h, with additional vortexing every 15min. Following centrifugation at 16,000xg for 15min to pellet protein and cell debris, the lipid containing liquid phase was transferred to a new tube. Extracts were respun to pellet remaining debris, then transferred to autosampler vials with insert for analysis.

### Untargeted lipidomics

Lipid extracts were analyzed as previously described6. Briefly, an Agilent 1290 Infinity II liquid chromatograph (LC) coupled to an Agilent 6546 quadrupole time-of-flight mass spectrometer (MS) was used for untargeted analyses. Lipids were separated using a Waters Acquity BEH C18 column (1.7μm, 2.1x100mm) held at 50°C. Mobile phase A was 60:40 MeCN:H2O and B was 9:1:90 MeCN:H2O:IPA, both with 0.1% (v/v) formic acid and 10mM ammonium formate, using the following gradient: 15% mobile phase B to 30% at 2.40min, to 48% at 3min, to 82% at 13.2min, finally to 99% from 13.2 to 13.8min held until 15.4 min before re-equilibration to 15% at a constant 0.500mL/min. Samples were analyzed in positive and negative ionization as separate experiments. All samples were injected undiluted at 10μl volumes for negative mode. For positive ionization, samples were diluted 20 to 40-fold (after evaluating saturation) and injected between 1-3uL.

Libraries of identified lipids were created using pooled samples and collecting MS/MS spectra on 6 consecutive injections using iterative exclusion. Agilent LipidAnnotator was used for lipid annotation, then individual samples were analyzed using MS1 acquisition and peak integration was performed in Agilent Profinder using mass and retention time from the lipid libraries. Subsequent steps including normalization to IS was performed in R using publicly available custom scripts (https://github.com/RJain52/Multi-Tissue-Cold-Exposure-Lipidomics).

### Targeted BMP and PG lipid measurements

A targeted BMP and PG LC/MS method was developed using an Agilent 1290 Infinity I LC coupled to an Agilent 6495c triple-quadrupole MS. Lipids were extracted using a one-phase extraction (see *Lipid extraction*)34,38. An *in-silico* multiple reaction monitoring (MRM) method was developed using LipidCreator for fragmentation patterns of BMP and PG of different acyl chain configurations in positive ([M+H]+ and [M+NH4]+ adducts) and negative ([M-H]-) ionization56. Samples were then injected using the same LC parameters as *Untargeted lipidomics* using the MRM methods. BMP lipids eluted 0.3-1min earlier than the PG species of the same acyl chains, depending on the species. This pattern was confirmed using BMP 18:1_18:1 and PG 18:1_18:1 standards purchased from Avanti Polar Lipids. Additionally, BMP(s) had unique fragments in positive ionization when compared to PG(s) of the same species. No unique fragments were observed in negative ionization; however the acyl chain carbon lengths were discernable, and ion signals were much higher due to the physicochemical properties of BMP and PG. Therefore, we created a negative mode dynamic MRM (dMRM) method for all identifiable BMP and PG species using retention time information from positive ionization, in which we could distinguish isomeric BMP and PG species via fragmentation, but scanning for precursor/products providing acyl chain information. For all subsequent analyses, pooled extracts were run in positive and negative ionization to allow for manual verification of lipid identity and individual samples were run only using negative mode dMRM analysis. Non-endogenous BMP 14:0/14:0 and PG 15:0/18:1d7 were used as IS. Data was processed in Agilent MassHunter Quantitative Analysis prior to downstream analyses in R.

### Non-esterified fatty acid measurements

Plasma non-esterified fatty acids were measured using a colorimetric assay on a Tecan plate reader according to manufacturer’s instructions. A five-point standard curve (0∼2.5 mM) of non-esterified oleic acid standard was used to calculate NEFA concentration from 2μl plasma. Samples were analyzed in duplicate.

### Ketone body assay

Plasma ketone bodies were measured by colorimetric assay using the manufacturer protocol. Briefly, 40μl plasma were diluted with 60μl assay buffer then duplicate wells for each sample were prepped with 50μl diluted plasma/well. Reaction was initiated and allowed to run for 30min at RT in the dark, then read using a Tecan plate reader at an absorbance of 445nm.

### Cathepsin B assay

Cathepsin B was assayed as previously described30. Briefly, 15mg liver was homogenized in 300μl lysis buffer (50mM Tris, 130mM NaCl, 10% (v/v) glycerol, 0.5% (v/v) NP-40, 0.5mM EDTA, 0.5mM EGTA, 1mM PMSF; pH=7.4) using sonication (Branson Sonifier 450) (60% duty cycle; output=4) on ice. Protein content was measured using BCA assay, then 15ug protein was incubated with 50uM Cathepsin B substrate. Reactions were conducted in the presence and absence of the cathepsin B inhibitor E-64-D in assay buffer (100mM NaCl, 100mM sodium acetate; pH=5.5) at a final volume of 100μl for 30min at 37°C. Activity was based on fluorescence of cleaved substrate measured at an excitation/emission of 355/460nm.

### Liposome assay

Liposome assays were carried out as previously described57. Briefly, S,S-(3,3’-diC18:1)-BMP/1,2-*O*-dioctadecenyl-*sn*-glycero-3-phosphocholine (DODPC, non-substrate of Pla2g15) liposomes were prepared by mixing reagents in a glass tube and drying down under N2. Lipid mixture was dispersed in a 50mM sodium acetate solution (pH 4.5) using sonication. To avoid the possibility of DFP affecting lipid membranes, DFP-treated Pla2g15 was dialyzed against 0.25 M sucrose/25 mM Hepes (pH 7.4)/1 mM EDTA before use. The UV spectrum of DFP-treated Pla2g15 after dialysis was compared to DFP-untreated Pla2g15 to confirm the structural properties of the protein. There was no significant difference in the spectra between the DFP-conjugated Pla2g15 and the untreated Pla2g15. Reactions contained liposomes and either sucrose buffer, sucrose buffer with human Pla2g15, or sucrose buffer with DFP-treated human Pla2g15, incubated at 37°C for 10min. Reactions were quenched with 2:1 chloroform:MeOH (2:1, v/v), then 0.9% (w/v) NaCl solution causing phase separation of a top aqueous and bottom hydrophobic layer. The bottom hydrophobic layer was transferred to glass tubes, dried down under N2, and developed using thin-layer chromatography with a solvent system of chloroform:MeOH:pyridine (99:1:2, v/v) to separate lipid fractions. Bands corresponding to lipid fractions were visualized by heating the plate treated with 8% (w/v) CuSO4 solution with 6.8% (v/v) H3PO4 and 32% (v/v) MeOH^57^.

### Statistical analyses

Analyses were conducted using R unless otherwise stated. Where relevant, mean±SD were reported. For comparisons between two groups, Student’s t-test was used and a *p*-value less than 0.05 was considered significant, unless otherwise stated. For untargeted analyses, false-discovery correction was used an *q*<0.05 was considered significant. For comparisons of >2 groups, ANOVA analysis was used to test for significance followed by pairwise t-tests. All scripts used for analysis are freely available on Github (username: RJain52).

## References

Abe, A., Hinkovska-Galcheva, V., Bouchev, P., Bouley, R., and Shayman, J.A. (2024). The role of lysosomal phospholipase A2 in the catabolism of bis(monoacylglycerol)phosphate and association with phospholipidosis. Journal of Lipid Research 65.

Abe, A., Hiraoka, M., and Shayman, J.A. (2006). Positional specificity of lysosomal phospholipase A2. J Lipid Res 47, 2268–2279.

Abe, A., Hiraoka, M., and Shayman, J.A. (2007). The acylation of lipophilic alcohols by lysosomal phospholipase A2. J Lipid Res 48, 2255–2263.

Abe, A., Kelly, R., Kollmeyer, J., Hiraoka, M., Lu, Y., and Shayman, J.A. (2008). The secretion and uptake of lysosomal phospholipase A2 by alveolar macrophages. J Immunol 181, 7873–7881.

Abu-Remaileh, M., Wyant, G.A., Kim, C., Laqtom, N.N., Abbasi, M., Chan, S.H., Freinkman, E., and Sabatini, D.M. (2017). Lysosomal metabolomics reveals V-ATPase- and mTOR-dependent regulation of amino acid efflux from lysosomes. Science (New York, NY) 358, 807–813.

Akgoc, Z., Sena-Esteves, M., Martin, D.R., Han, X., d’Azzo, A., and Seyfried, T.N. (2015). Bis(monoacylglycero)phosphate: a secondary storage lipid in the gangliosidoses. J Lipid Res 56, 1006–1013.

Akira, A., Miki, H., and James, A.S. (2007). A Role for Lysosomal Phospholipase A2 in Drug Induced Phospholipidosis. Drug Metabolism Letters 1, 49–53.

Alvarez-Valadez, K., Sauvat, A., Diharce, J., Leduc, M., Stoll, G., Guittat, L., Lambertucci, F., Paillet, J., Motiño, O., Ferret, L., et al. (2024). Lysosomal damage due to cholesterol accumulation triggers immunogenic cell death. Autophagy, 1-23.

Anderson, D.M.G., Ablonczy, Z., Koutalos, Y., Hanneken, A.M., Spraggins, J.M., Calcutt, M.W., Crouch, R.K., Caprioli, R.M., and Schey, K.L. (2017). Bis(monoacylglycero)phosphate lipids in the retinal pigment epithelium implicate lysosomal/endosomal dysfunction in a model of Stargardt disease and human retinas. Sci Rep 7, 17352.

Azimifar, S.B., Nagaraj, N., Cox, J., and Mann, M. (2014). Cell-type-resolved quantitative proteomics of murine liver. Cell Metab 20, 1076–1087.

Body, D.R., and Gray, G.M. (1967). The isolation and characterisation of phosphatidylglycerol and a structural isomer from pig lung. Chemistry and Physics of Lipids 1, 254–263.

Boland, S., Swarup, S., Ambaw, Y.A., Malia, P.C., Richards, R.C., Fischer, A.W., Singh, S., Aggarwal, G., Spina, S., Nana, A.L., et al. (2022). Deficiency of the frontotemporal dementia gene GRN results in gangliosidosis. Nat Commun 13, 5924.

Bornstein, M.R., Neinast, M.D., Zeng, X., Chu, Q., Axsom, J., Thorsheim, C., Li, K., Blair, M.C., Rabinowitz, J.D., and Arany, Z. (2023). Comprehensive quantification of metabolic flux during acute cold stress in mice. Cell Metab 35, 2077–2092.e2076.

Bouley, R.A., Hinkovska-Galcheva, V., Shayman, J.A., and Tesmer, J.J.G. (2019). Structural Basis of Lysosomal Phospholipase A2 Inhibition by Zn2+. Biochemistry 58, 1709–1717.

Bulfon, D., Breithofer, J., Grabner, G.F., Fawzy, N., Pirchheim, A., Wolinski, H., Kolb, D., Hartig, L., Tischitz, M., Zitta, C., et al. (2024). Functionally overlapping intra- and extralysosomal pathways promote bis(monoacylglycero)phosphate synthesis in mammalian cells. Nature Communications 15, 9937.

Burke, J.E., and Dennis, E.A. (2009). Phospholipase A2 structure/function, mechanism, and signaling. J Lipid Res 50 *Suppl*, S237-242.

Chen, J., Cazenave-Gassiot, A., Xu, Y., Piroli, P., Hwang, R., DeFreitas, L., Chan, R.B., Di Paolo, G., Nandakumar, R., Wenk, M.R., et al. (2023). Lysosomal phospholipase A2 contributes to the biosynthesis of the atypical late endosome lipid bis(monoacylglycero)phosphate. Communications Biology 6, 210.

Chen, S., Francioli, L.C., Goodrich, J.K., Collins, R.L., Kanai, M., Wang, Q., Alföldi, J., Watts, N.A., Vittal, C., Gauthier, L.D., et al. (2024). A genomic mutational constraint map using variation in 76,156 human genomes. Nature 625, 92–100.

Chitraju, C., Fischer, A.W., Farese, R.V., Jr., and Walther, T.C. (2020). Lipid Droplets in Brown Adipose Tissue Are Dispensable for Cold-Induced Thermogenesis. Cell reports 33, 108348.

Eapen, V.V., Swarup, S., Hoyer, M.J., Paulo, J.A., and Harper, J.W. (2021). Quantitative proteomics reveals the selectivity of ubiquitin-binding autophagy receptors in the turnover of damaged lysosomes by lysophagy. Elife 10.

Glukhova, A., Hinkovska-Galcheva, V., Kelly, R., Abe, A., Shayman, J.A., and Tesmer, J.J.G. (2015). Structure and function of lysosomal phospholipase A2 and lecithin:cholesterol acyltransferase. Nat Commun 6, 6250.

Gosis, B.S., Wada, S., Thorsheim, C., Li, K., Jung, S., Rhoades, J.H., Yang, Y., Brandimarto, J., Li, L., Uehara, K., et al. (2022). Inhibition of nonalcoholic fatty liver disease in mice by selective inhibition of mTORC1. Science (New York, NY) 376, eabf8271.

Grabner, G.F., Fawzy, N., Pribasnig, M.A., Trieb, M., Taschler, U., Holzer, M., Schweiger, M., Wolinski, H., Kolb, D., Horvath, A., et al. (2019). Metabolic disease and ABHD6 alter the circulating bis(monoacylglycerol)phosphate profile in mice and humans. J Lipid Res 60, 1020–1031.

Grabner, G.F., Fawzy, N., Schreiber, R., Pusch, L.M., Bulfon, D., Koefeler, H., Eichmann, T.O., Lass, A., Schweiger, M., Marsche, G., et al. (2020). Metabolic regulation of the lysosomal cofactor bis(monoacylglycero)phosphate in mice. Journal of lipid research 61, 995–1003.

Haemmerle, G., Lass, A., Zimmermann, R., Gorkiewicz, G., Meyer, C., Rozman, J., Heldmaier, G., Maier, R., Theussl, C., Eder, S., et al. (2006). Defective lipolysis and altered energy metabolism in mice lacking adipose triglyceride lipase. Science 312, 734–737.

Hinkovska-Galcheva, V., Kelly, R., Manthei, K.A., Bouley, R., Yuan, W., Schwendeman, A., Tesmer, J.J.G., and Shayman, J.A. (2018). Determinants of pH profile and acyl chain selectivity in lysosomal phospholipase A(2). J Lipid Res 59, 1205–1218.

Hiraoka, M., Abe, A., Lu, Y., Yang, K., Han, X., Gross, R.W., and Shayman, J.A. (2006). Lysosomal phospholipase A2 and phospholipidosis. Molecular and cellular biology 26, 6139–6148.

Ilnytska, O., Lai, K., Gorshkov, K., Schultz, M.L., Tran, B.N., Jeziorek, M., Kunkel, T.J., Azaria, R.D., McLoughlin, H.S., Waghalter, M., et al. (2021). Enrichment of NPC1-deficient cells with the lipid LBPA stimulates autophagy, improves lysosomal function, and reduces cholesterol storage. The Journal of biological chemistry 297, 100813.

Jain, R., Davidson, J., Gonzalez, P., Coe, C., King, C., Ryff, C., Bersh, A., Mohsin, S., Love, G.D., Nimityongskul, F., et al. (2022). Identification of novel plasma lipid markers of cardiovascular disease risk in White and Black women. medRxiv, 2022.2008.2024.22279186.

Jurado-Fasoli, L., Sanchez-Delgado, G., Di, X., Yang, W., Kohler, I., Villarroya, F., Aguilera, C.M., Hankemeier, T., Ruiz, J.R., and Martinez-Tellez, B. (2024). Cold-induced changes in plasma signaling lipids are associated with a healthier cardiometabolic profile independently of brown adipose tissue. Cell Rep Med 5, 101387.

Kagohashi, Y., Sasaki, M., May, A.I., Kawamata, T., and Ohsumi, Y. (2023). The mechanism of Atg15-mediated membrane disruption in autophagy. J Cell Biol 222.

Kirkegaard, T., Roth, A.G., Petersen, N.H., Mahalka, A.K., Olsen, O.D., Moilanen, I., Zylicz, A., Knudsen, J., Sandhoff, K., Arenz, C., et al. (2010). Hsp70 stabilizes lysosomes and reverts Niemann-Pick disease-associated lysosomal pathology. Nature 463, 549–553.

Laqtom, N.N., Dong, W., Medoh, U.N., Cangelosi, A.L., Dharamdasani, V., Chan, S.H., Kunchok, T., Lewis, C.A., Heinze, I., Tang, R., et al. (2022). CLN3 is required for the clearance of glycerophosphodiesters from lysosomes. Nature 609, 1005–1011.

Li, Y., Wang, X., Li, M., Yang, C., and Wang, X. (2022). M05B5.4 (lysosomal phospholipase A2) promotes disintegration of autophagic vesicles to maintain C. elegans development. Autophagy 18, 595–607.

Locatelli-Hoops, S., Remmel, N., Klingenstein, R., Breiden, B., Rossocha, M., Schoeniger, M., Koenigs, C., Saenger, W., and Sandhoff, K. (2006). Saposin A Mobilizes Lipids from Low Cholesterol and High Bis(monoacylglycerol)phosphate-containing Membranes: PATIENT VARIANT SAPOSIN A LACKS LIPID EXTRACTION CAPACITY*. Journal of Biological Chemistry 281, 32451–32460.

Logan, T., Simon, M.J., Rana, A., Cherf, G.M., Srivastava, A., Davis, S.S., Low, R.L.Y., Chiu, C.L., Fang, M., Huang, F., et al. (2021). Rescue of a lysosomal storage disorder caused by Grn loss of function with a brain penetrant progranulin biologic. Cell 184, 4651–4668.e4625.

Longato, L., Tong, M., Wands, J.R., and de la Monte, S.M. (2012). High fat diet induced hepatic steatosis and insulin resistance: Role of dysregulated ceramide metabolism. Hepatol Res 42, 412–427.

Medoh, U.N., and Abu-Remaileh, M. (2024). The Bis(monoacylglycero)-phosphate Hypothesis: From Lysosomal Function to Therapeutic Avenues. Annu Rev Biochem 93, 447–469.

Medoh, U.N., Hims, A., Chen, J.Y., Ghoochani, A., Nyame, K., Dong, W., and Abu-Remaileh, M. (2023). The Batten disease gene product CLN5 is the lysosomal bis(monoacylglycero)phosphate synthase. Science 381, 1182–1189.

Niu, L., Geyer, P.E., Gupta, R., Santos, A., Meier, F., Doll, S., Wewer Albrechtsen, N.J., Klein, S., Ortiz, C., Uschner, F.E., et al. (2022). Dynamic human liver proteome atlas reveals functional insights into disease pathways. Molecular Systems Biology 18, e10947.

Nyame, K., Hims, A., Aburous, A., Laqtom, N.N., Dong, W., Medoh, U.N., Heiby, J.C., Xiong, J., Ori, A., and Abu-Remaileh, M. (2024a). Glycerophosphodiesters inhibit lysosomal phospholipid catabolism in Batten disease. Mol Cell 84, 1354–1364.e1359.

Nyame, K., Xiong, J., de Jong, A.P., Alsohybe, H.N., Raaben, M., Hartmann, G., Simcox, J.A., Blomen, V.A., and Abu-Remaileh, M. (2024b). PLA2G15 is a Lysosomal BMP Hydrolase with Ester Position Specificity and its Targeting Ameliorates Lysosomal Disease. bioRxiv.

Oninla, V.O., Breiden, B., Babalola, J.O., and Sandhoff, K. (2014). Acid sphingomyelinase activity is regulated by membrane lipids and facilitates cholesterol transfer by NPC2. J Lipid Res 55, 2606–2619.

Palmieri, M., Impey, S., Kang, H., di Ronza, A., Pelz, C., Sardiello, M., and Ballabio, A. (2011). Characterization of the CLEAR network reveals an integrated control of cellular clearance pathways. Human Molecular Genetics 20, 3852–3866.

Rodriguez-Navarro, J.A., Kaushik, S., Koga, H., Dall’Armi, C., Shui, G., Wenk, M.R., Di Paolo, G., and Cuervo, A.M. (2012). Inhibitory effect of dietary lipids on chaperone-mediated autophagy. Proc Natl Acad Sci U S A 109, E705–714.

Sardiello, M., Palmieri, M., di Ronza, A., Medina, D.L., Valenza, M., Gennarino, V.A., Di Malta, C., Donaudy, F., Embrione, V., Polishchuk, R.S., et al. (2009). A gene network regulating lysosomal biogenesis and function. Science (New York, NY) 325, 473–477.

Scherer, M., Schmitz, G., and Liebisch, G. (2010). Simultaneous Quantification of Cardiolipin, Bis(monoacylglycero)phosphate and their Precursors by Hydrophilic Interaction LC−MS/MS Including Correction of Isotopic Overlap. Analytical Chemistry 82, 8794–8799.

Schreiber, R., Diwoky, C., Schoiswohl, G., Feiler, U., Wongsiriroj, N., Abdellatif, M., Kolb, D., Hoeks, J., Kershaw, E.E., Sedej, S., et al. (2017). Cold-Induced Thermogenesis Depends on ATGL-Mediated Lipolysis in Cardiac Muscle, but Not Brown Adipose Tissue. Cell Metab 26, 753–763.e757.

Settembre, C., De Cegli, R., Mansueto, G., Saha, P.K., Vetrini, F., Visvikis, O., Huynh, T., Carissimo, A., Palmer, D., Jürgen Klisch, T., et al. (2013). TFEB controls cellular lipid metabolism through a starvation-induced autoregulatory loop. Nature cell biology 15, 647–658.

Severgnini, M., Sherman, J., Sehgal, A., Jayaprakash, N.K., Aubin, J., Wang, G., Zhang, L., Peng, C.G., Yucius, K., Butler, J., et al. (2012). A rapid two-step method for isolation of functional primary mouse hepatocytes: cell characterization and asialoglycoprotein receptor based assay development. Cytotechnology 64, 187–195.

Shayman, J.A., and Tesmer, J.J.G. (2019). Lysosomal phospholipase A2. Biochim Biophys Acta Mol Cell Biol Lipids 1864, 932–940.

Showalter, M.R., Berg, A.L., Nagourney, A., Heil, H., Carraway, K.L., 3rd, and Fiehn, O. (2020). The Emerging and Diverse Roles of Bis(monoacylglycero) Phosphate Lipids in Cellular Physiology and Disease. Int J Mol Sci *21*.

Simcox, J., Geoghegan, G., Maschek, J.A., Bensard, C.L., Pasquali, M., Miao, R., Lee, S., Jiang, L., Huck, I., Kershaw, E.E., et al. (2017). Global Analysis of Plasma Lipids Identifies Liver-Derived Acylcarnitines as a Fuel Source for Brown Fat Thermogenesis. Cell Metab 26, 509–522.e506.

Singh, S., Dransfeld, U.E., Ambaw, Y.A., Lopez-Scarim, J., Farese, R.V., and Walther, T.C. (2024). PLD3 and PLD4 synthesize S,S-BMP, a key phospholipid enabling lipid degradation in lysosomes. Cell 187, 6820–6834.e6824.

Sostre-Colón, J., Uehara, K., Garcia Whitlock, A.E., Gavin, M.J., Ishibashi, J., Potthoff, M.J., Seale, P., and Titchenell, P.M. (2021). Hepatic AKT orchestrates adipose tissue thermogenesis via FGF21-dependent and -independent mechanisms. Cell Rep 35, 109128.

Taniyama, Y., Shibata, S., Kita, S., Horikoshi, K., Fuse, H., Shirafuji, H., Sumino, Y., and Fujino, M. (1999). Cloning and expression of a novel lysophospholipase which structurally resembles lecithin cholesterol acyltransferase. Biochem Biophys Res Commun 257, 50–56.

Varadharajan, V., Ramachandiran, I., Massey, W.J., Jain, R., Banerjee, R., Horak, A.J., McMullen, M.R., Huang, E., Bellar, A., Lorkowski, S.W., et al. (2024). Membrane-bound O-acyltransferase 7 (MBOAT7) shapes lysosomal lipid homeostasis and function to control alcohol-associated liver injury. eLife 12.

Xia, J.Y., Holland, W.L., Kusminski, C.M., Sun, K., Sharma, A.X., Pearson, M.J., Sifuentes, A.J., McDonald, J.G., Gordillo, R., and Scherer, P.E. (2015). Targeted Induction of Ceramide Degradation Leads to Improved Systemic Metabolism and Reduced Hepatic Steatosis. Cell Metab 22, 266–278.

Yu, Y., Gao, S.M., Guan, Y., Hu, P.-W., Zhang, Q., Liu, J., Jing, B., Zhao, Q., Sabatini, D.M., Abu-Remaileh, M., et al. (2024). Organelle proteomic profiling reveals lysosomal heterogeneity in association with longevity. eLife 13, e85214.

Zhang, S., Williams, K.J., Verlande-Ferrero, A., Chan, A.P., Su, G.B., Kershaw, E.E., Cox, J.E., Maschek, J.A., Shapira, S.N., Christofk, H.R., et al. (2024). Acute activation of adipocyte lipolysis reveals dynamic lipid remodeling of the hepatic lipidome. J Lipid Res 65, 100434.

