## Supplemental figures for "Modulation of hepatic transcription factor EB activity during cold exposure uncovers direct regulation of bis(monoacylglycero)phosphate lipids by *Pla2g15*"

### Supplementary Figure 1

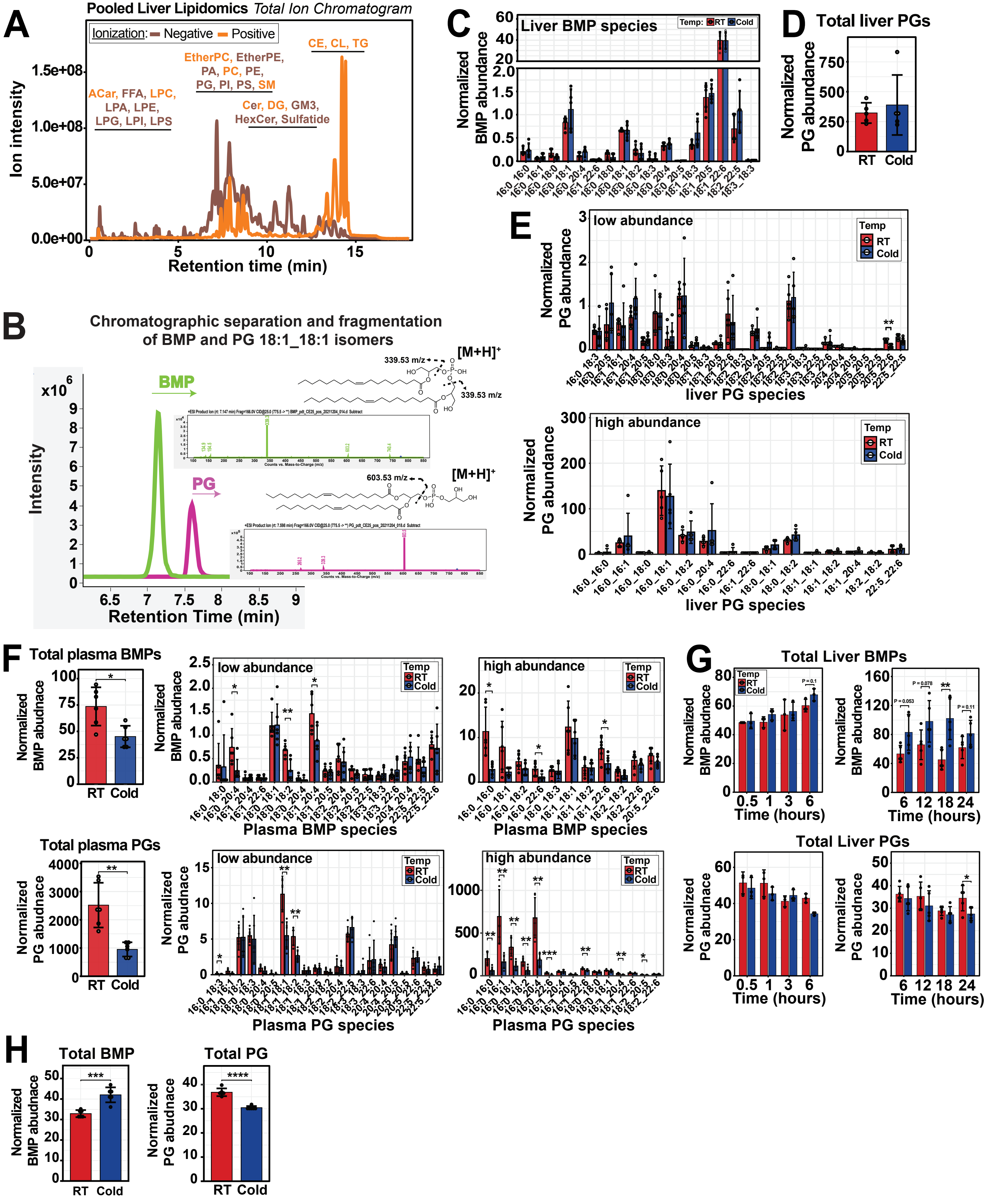

### Supplementary Figure 2

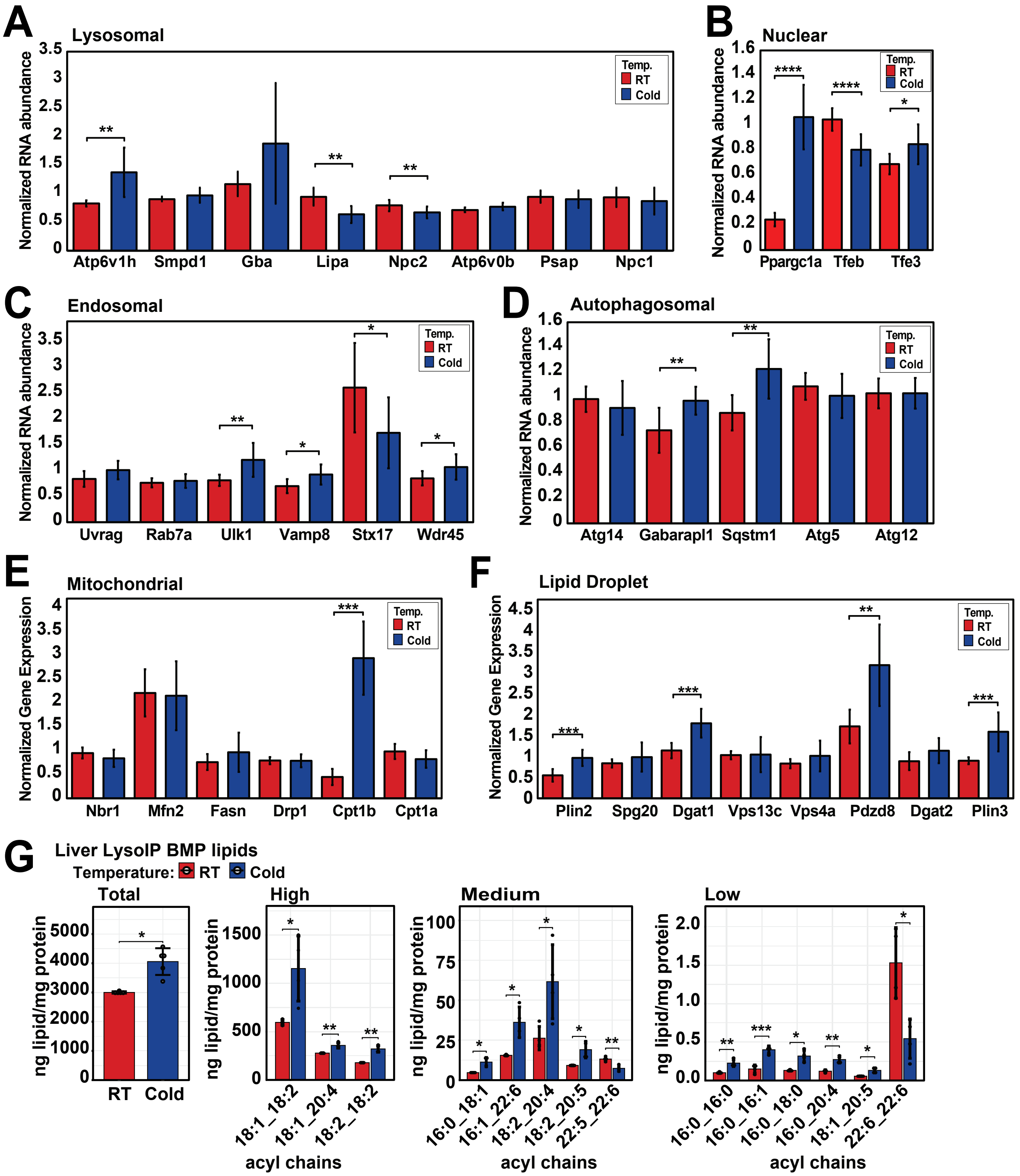

### Supplementary Figure 3

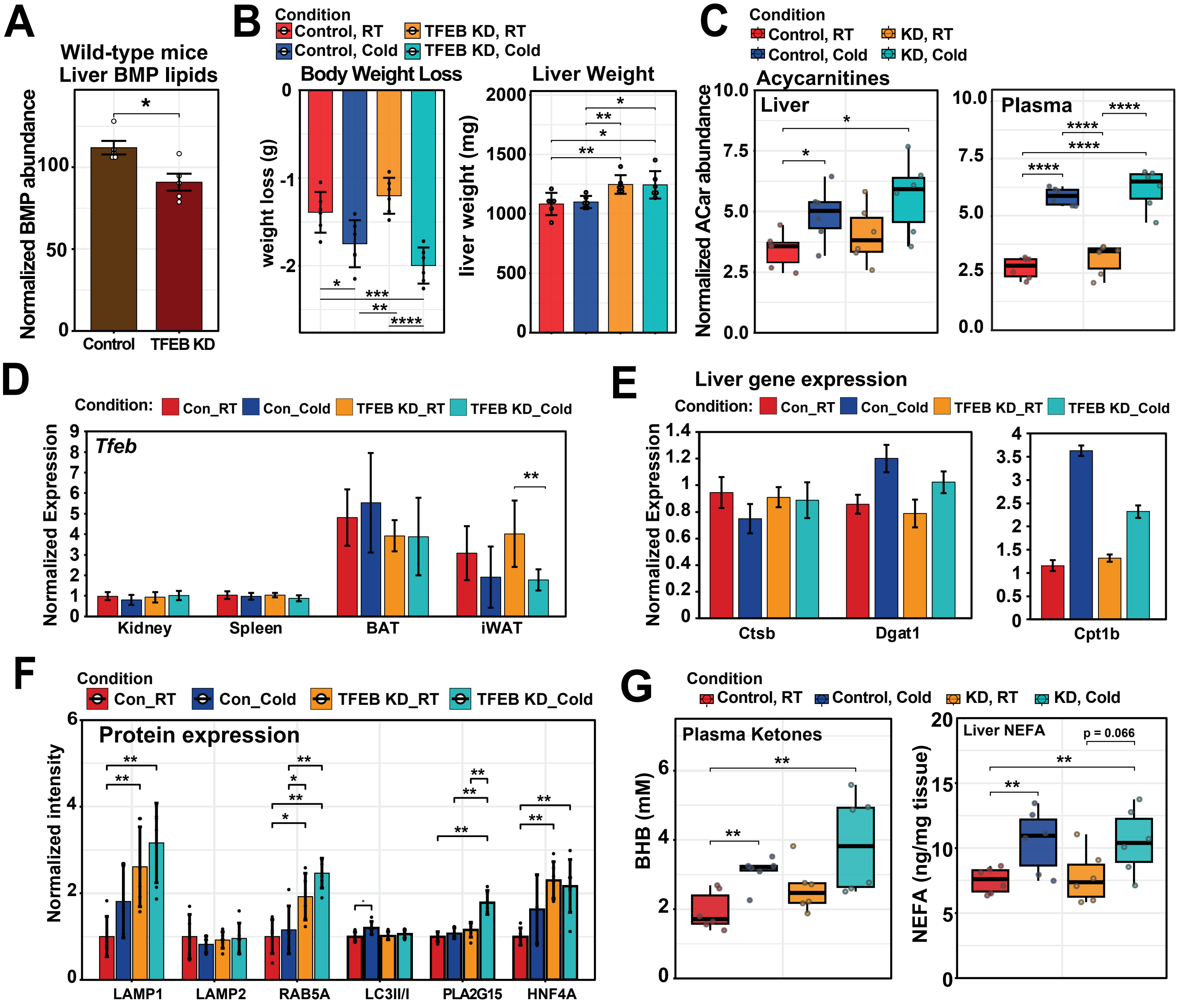

### Supplementary Figure 4

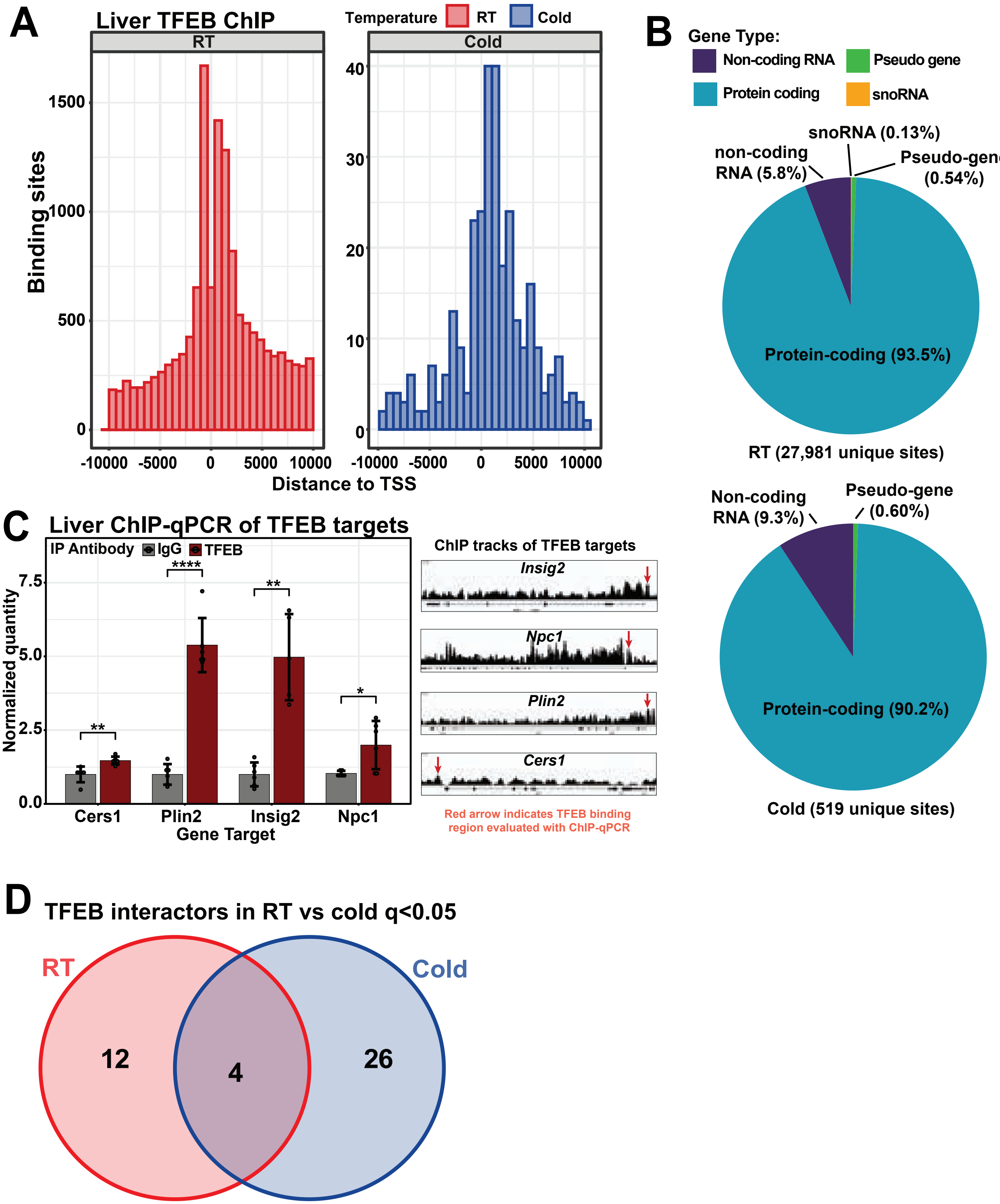

### Supplementary Figure 5

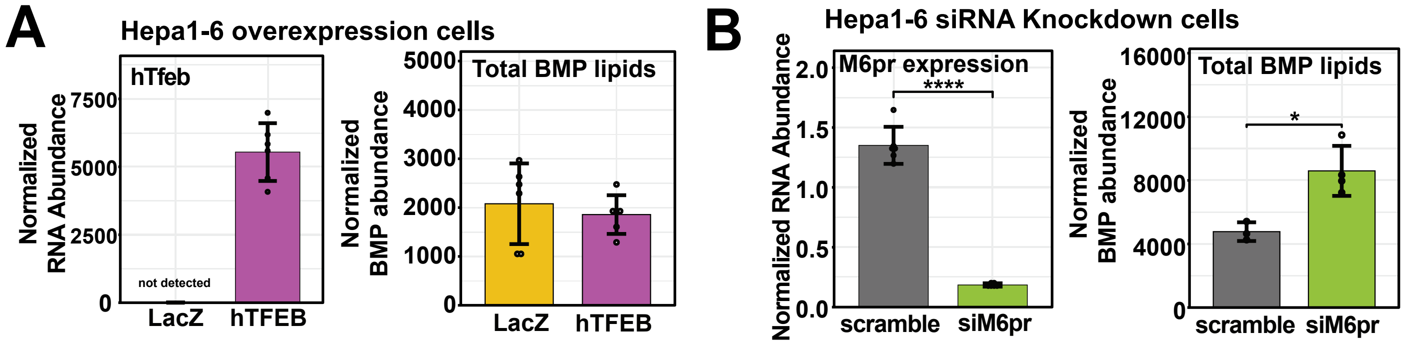

### Supplementary Figure 6

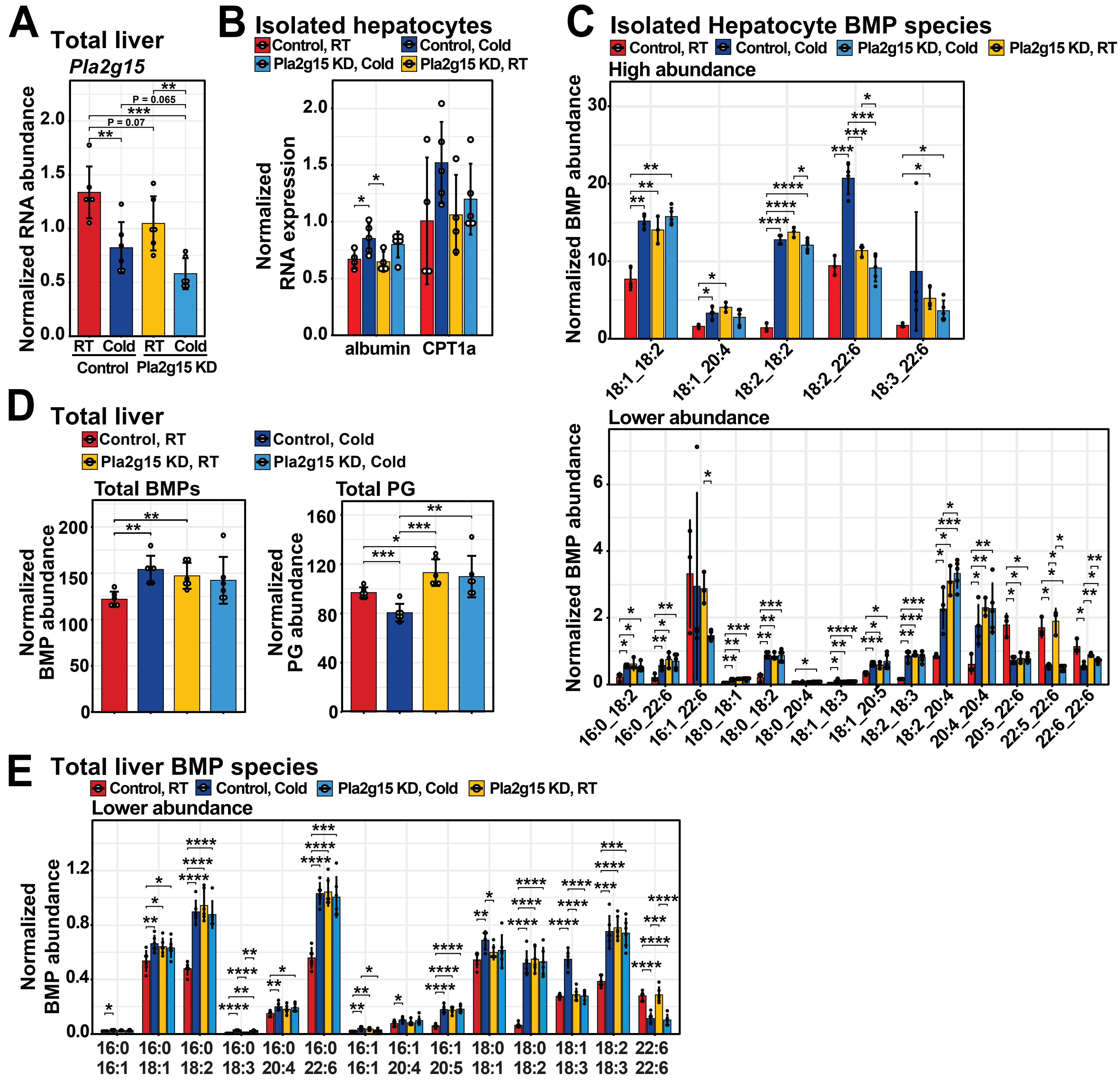

### Supplementary Figure 7

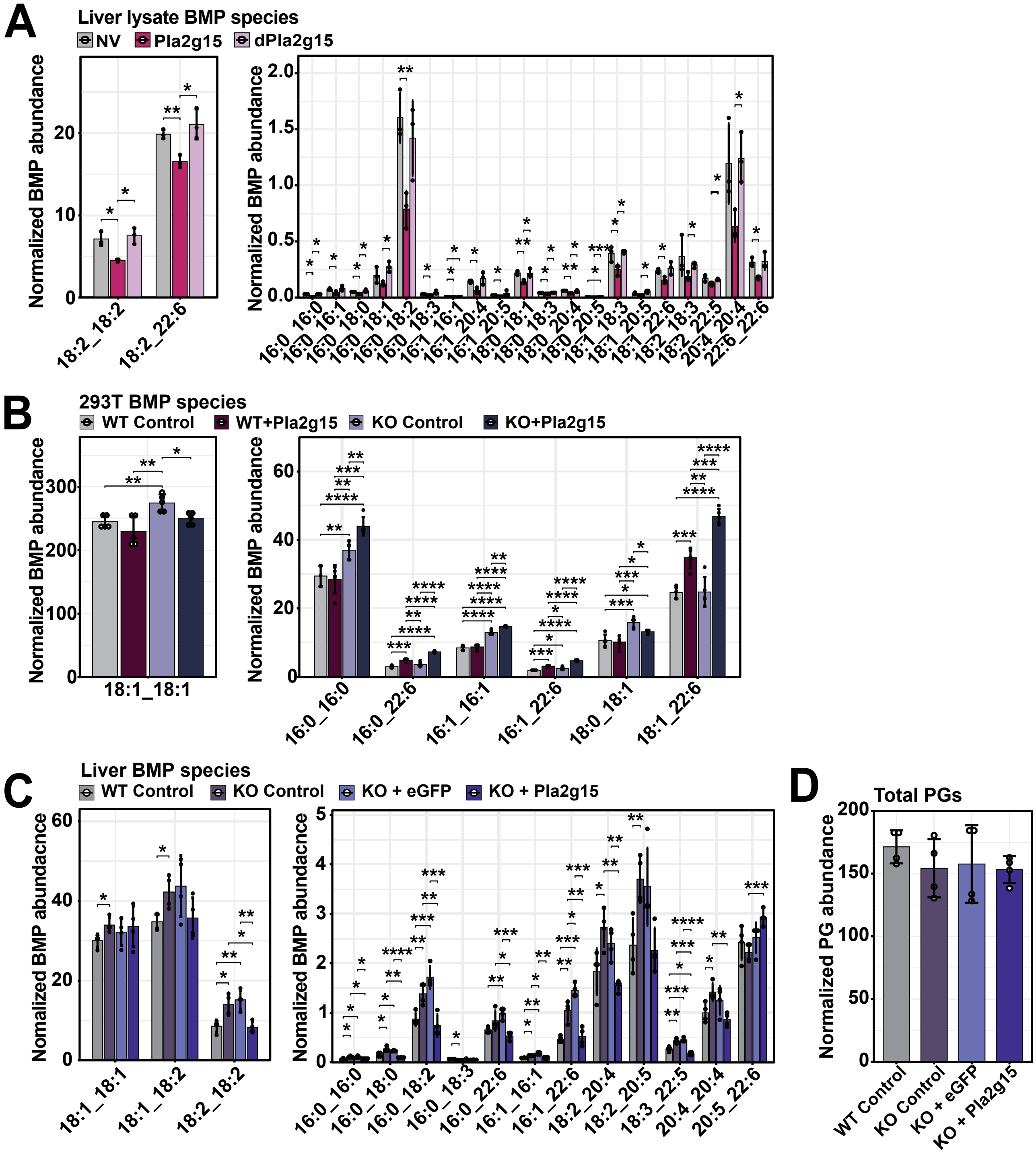
